# A transcriptional network required for *Toxoplasma gondii* tissue cyst formation is dispensable for long-term persistence

**DOI:** 10.1101/2022.04.06.487408

**Authors:** Sarah Sokol Borrelli, Sarah M. Reilly, Katherine G. Sharp, Leah F. Cabo, Hisham S. Alrubaye, Bruno Martorelli Di Genova, Jon P. Boyle

## Abstract

Cyst formation is a key feature of the *T. gondii* life cycle but the genetic networks that drive this process are not yet fully characterized. To identify new components of this network, we compared *T. gondii* to its nearest extant relative *Hammondia hammondi* given the critical differences between these species in the timing and efficiency of cyst formation. Using transcriptional data from critical developmental and pH exposure time points from both species, we identified the gene TGVEG_311100, which we named *Regulator of Cystogenesis 1* (*ROCY1*), as being both necessary and sufficient for cyst formation in *T. gondii*. Compared to WT parasites, TGVEGΔROCY1 parasites formed significantly fewer tissue cysts in response to alkaline pH stress *in vitro* and cysts were nearly undetectable in mouse brains for up to 9 weeks post-infection. Overexpression of tagged ROCY1 in WT parasites was sufficient to induce cyst formation *in vitro* in both WT and ROCY1-deficient parasites, demonstrating that ROCY1 is both necessary and sufficient for cyst formation. Moreover this induction of cyst formation required at least 1 of 3 predicted CCCH Zinc finger domains. Mice chronically infected with ΔROCY1 parasites had detectable tachyzoites in the brain for up to 37 days post-infection (while mice infected with WT parasites did not), and CNS transcriptional analyses at day 30 post-infection throughout the chronic phase of infection revealed inflammatory signatures consistent with acute infection in ΔROCY1 parasites compared to WT. Despite our inability to detect brain cysts in infected mice, both WT and ΔROCY1 knockout parasites reactivated after dexamethasone treatment with similar timing and magnitude for up to 5 months post infection, challenging the paradigm that long term parasite persistence in the CNS requires cyst formation. These data identify a new regulator of cyst formation in *T. gondii* that is both necessary and sufficient for cyst formation, and whose function relies on its conserved nucleic acid binding motif.

## Introduction

Eukaryotic parasites are renowned for their complex, multihost life cycles that require transitions between life stages that differ dramatically in their morphology, metabolism and reproductive niche(s). *Toxoplasma gondii* is no exception, a tissue-dwelling coccidian that has infected a third of the world’s population. A renowned feature of the *T. gondii* life cycle is the ability to persist for years as a bradyzoite, a slow-growing, intracellular life stage surrounded by a sugar-rich cyst wall. Despite their importance for transmission and disease, a number of genetic components used by *T. gondii* to transition between life stages have been discovered (Farhat *et al*., 2020; Huang *et al*., 2017; Radke *et al*., 2013, 2018; Waldman *et al*., 2020), but to date their complete regulatory networks are unknown as are the mechanisms that lead to their activation. It has long been assumed that tissue cyst formation is a response to immune pressure within the host and is required for *T. gondii* persistence in the host (Kim *et al*., 2007; Tomita *et al*., 2013). In mice the primary site of persistence is within the brain where it resides within neurons and can cause lethal encephalitis when immune surveillance is altered by experimental treatment or disease (Cabral *et al*., 2020; Jones *et al*., 1996; Suzuki *et al*., 1989; Takashima *et al*., 2008). A less than complete understanding of how this process is regulated transcriptionally and executed by downstream effectors presents a barrier to the development of new therapies targeting this drug-refractory life stage and limits our ability to understand the importance of cyst formation itself in *T. gondii* infection, persistence and transmission to definitive or intermediate hosts.

As a means to identify additional regulators of cyst formation in T *gondii*, we have taken advantage of the *T. gondii*/*H. hammondi* comparative system. *H. hammondi* is the nearest extant relative of *T. gondii* and shared nearly all of its genes across chromosomes that are almost entirely syntenic (Dubey and Sreekumar, 2003; Lorenzi *et al*., 2016; Walzer *et al*., 2013, 2014). These two parasite species are morphologically and antigenically similar, share the same definitive host and organelles, and transition between similar life stages as they progress through their life cycle (Dubey and Ferguson, 2015; Dubey and Sreekumar, 2003; Frenkel and Dubey, 1975). Even though these parasites have many similarities, *H. hammondi* follows a strictly obligate heteroxenous life cycle (Dubey and Ferguson, 2015; Frenkel and Dubey, 1975) while *T. gondii* has a facultative heteroxenous/homoxenous life cycle. This is driven by the unique ability of *T. gondii* bradyzoites, the slow-growing tissue cyst forming life stage of the parasite responsible for chronic infection, to convert back to tachyzoites, the rapidly replicating life stage that can lead to disease (Dubey, 1998, 2009). This unique interconversion phenomenon is not shared with *H. hammondi*. Bradyzoite to tachyzoite interconversion allows for *T. gondii* to pass between intermediate hosts, thus expanding its transmission dynamics (Dubey *et al*., 1998). Furthermore, this interconversion can be lethal or result in severe disease in immune compromised organ transplant patients, cancer patients, and individuals with HIV/AIDS (Derouin *et al*., 1987, 2008). The similarities between *T. gondii* and *H. hammondi* bradyzoites paired with the stark contrast in when and how bradyzoites form in each species makes interspecies comparison a promising strategy for uncovering critical components driving the bradyzoite developmental program. *H. hammondi’s* strict obligate heteroxenous life cycle restricts its ability to initiate infection in organisms other than its definitive feline host after it transitions into its bradyzoite life form (Dubey and Sreekumar, 2003; Riahi *et al*., 1995; Sheffield *et al*., 1976; Sokol *et al*., 2018). Moreover, *H. hammondi* cannot be induced to form bradyzoites in its early life stages when treated with alkaline pH stress (Sokol *et al*., 2018), a known and potent inducer of cystogenesis in *T. gondii* (Soête *et al*., 1994; Weiss *et al*., 1995, 1998). Yet, *H. hammondi* predictably and reliably forms tissue cysts spontaneously which is also accompanied by robust transcriptional changes resembling those observed in *T. gondii* as they differentiate into bradyzoites (Dubey and Sreekumar, 2003; Sokol *et al*., 2018).

Numerous studies designed to characterize the *T. gondii* bradyzoite have led to the identification of several bradyzoite specific proteins along with their function (Jeffers *et al*., 2018) and have characterized the major transcriptional differences that occur during bradyzoite development in *T. gondii* (Buchholz *et al*., 2011; Chen *et al*., 2018; Croken *et al*., 2014; Radke *et al*., 2005; Sharma *et al*., 2020; Tanaka *et al*., 2013). However, until recently the mechanisms used by *T. gondii* to initiate bradyzoite development were elusive. The first major class of transcription factors identified in Apicomplexans were AP2 transcription factors (Balaji *et al*., 2005), which begun to broaden our understanding of the regulation of bradyzoite development. These transcription factors play fundamental roles in regulating stage conversion associated gene expression in closely related *Plasmodium* species (Kafsack *et al*., 2014; Painter *et al*., 2011). In *T. gondii*, this family of proteins are important for cystogenesis and function as activators and repressors of stage conversion associated gene expression (Hong *et al*., 2017; Huang *et al*., 2017; Radke *et al*., 2013, 2018; Srivastava *et al*., 2020; Walker *et al*., 2013). In addition to AP2 factors, another *T. gondii* transcription factor that regulates the tachyzoite to bradyzoite transition, Bradyzoite-Formation Deficient-1 (BFD1) was recently identified as a transcription factor that completely abolishes tissue cyst formation *in vitro* and *in vivo* when depleted (Waldman *et al*., 2020). The identification of BFD1 as a regulator of differentiation was a fundamental finding and has greatly contributed to our understanding of the bradyzoite developmental pathway in *T. gondii*. However, our understanding of this process and all its major contributors remains unclear.

Here, we used a novel interspecies comparative approach that exploits our knowledge of the critical differences in tissue cyst formation between *T. gondii* and *H. hammondi* to identify factors that are important for driving cystogenesis. Using this system, we identified a *T. gondii* gene, *Regulator of Cystogenesis 1* (*ROCY1*), as a gene that is necessary and sufficient for cyst formation *in vitro* and permitted us to determine for the first time the precise role played by cyst formation itself in *T. gondii* persistence.

## Results

### Comparative transcriptomics between *T. gondii* and *H. hammondi* identifies candidate genes involved in tissue cyst development

To identify candidate *T. gondii* genes implicated in tissue cyst development, we leveraged species-specific differences in spontaneous and alkaline pH-induced tissue cyst formation that we previously identified in *T. gondii* and *H. hammondi* parasites (Sokol *et al*., 2018). When *T. gondii* sporozoites (VEG) are exposed to alkaline pH-induced stress conditions at 3 days post excystation (DPE), they robustly form Dolichos biflorus agglutinin (DBA)-positive tissue cysts *in vitro*. *H. hammondi* fails to form DBA-positive tissue cysts under these conditions, but spontaneously forms DBA-positive tissue cysts beginning at 12DPE (Summarized in Figure 1A) (Sokol *et al*., 2018).

**Figure 1.**
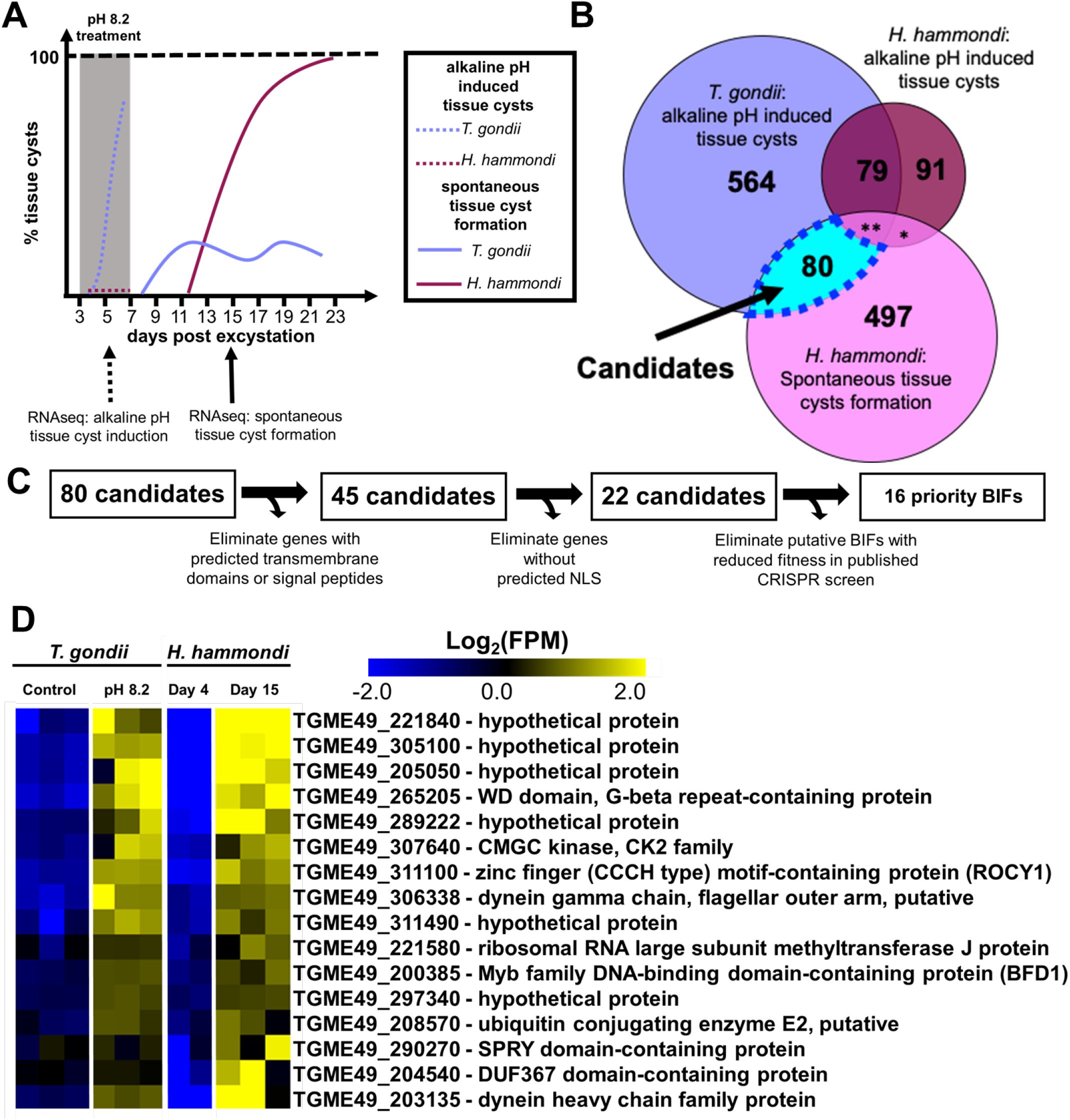
Comparative transcriptomics between *T. gondii* and *H. hammondi* identifies candidate genes involved in tissue cyst development. A) Schematic depicting spontaneous and alkaline pH induced tissue cyst formation phenotype over a 23-day time course in *T. gondii* VEG and *H. hammondi* Amer. B) Venn diagram showing genes with significant (|Log_2_ Fold Change| ≥ 1, P_adj_ < 0.01) differences in transcript abundance during alkaline pH induced stress in *H. hammondi* and *T. gondii* (at least 1 of 2 independent experiments) and during spontaneous development in *H. hammondi*. *=7, **=22 C) Schematic representing the prioritization strategy used to identify candidates predicted to be involved in tissue cyst formation. D) Mean centered heat map showing 16 priority candidate BIFs. Data from *T. gondii* and *H. hammondi* are each mean centered separately.

We hypothesized that some of the genes that change in transcript abundance during alkaline pH stress exposure in *T. gondii* and during spontaneous development in *H. hammondi* would have critical regulatory roles in tissue cyst development. To identify candidate genes that could be driving tissue cyst formation, we performed RNA sequencing experiments on *T. gondii* and *H. hammondi* sporozoite-derived infections exposed to alkaline pH, low serum, and CO_2_ starvation for 48 hours beginning at 3DPE and on *T. gondii* and *H. hammondi* grown in control conditions. We then used this new transcriptional data in combination with our previously published data (Sokol *et al*., 2018) to identify genes that had 1) significant changes in transcript abundance (|Log_2_FC| > 1, P_adj_ < 0.01) in *T. gondii* in alkaline pH stress conditions compared to control condition in at least 1 of 2 experiments, 2) genes that did not have significant changes (|Log_2_FC| < 1 and/or P_adj_ > 0.01) in *H. hammondi* during stressed conditions relative to control conditions in at least one of 2 experiments as *H. hammondi* fails to form DBA-positive tissue cysts in response to this treatment, and 3) genes with significant changes in transcript abundance (|Log_2_FC| > 1, P_adj_ < 0.01) during spontaneous tissue cyst development in *H. hammondi* (D4 vs D15). We identified 80 candidate genes fitting these criteria (Figure 1A-B). To prioritize these 80 candidates for further investigation, we sought to identify potential transcription factors that could be responsible for driving the global transcriptional changes needed for the transition from tachyzoite to bradyzoite. To do this we first removed candidates with a predicted signal peptide or transmembrane domains, leaving 48 candidates. Next, we identified 22 genes containing predicted nuclear localization sequences using NLSmapper (Kosugi *et al*., 2009) and NLStradamus (Nguyen Ba *et al*., 2009) using default settings. Finally, to further prioritize our candidate genes, we utilized data from a genome-wide CRISPR screen in *T. gondii* that predicts whether a gene contributes to tachyzoite fitness by calculating a CRISPR phenotype score from the ratio of the abundance of gRNAs present following *in vitro* growth in tachyzoite conditions to the abundance of gRNAs present in the initial library (Sidik *et al*., 2016). We then selected bradyzoite induction factors (BIFs) if their transcript abundance increased during tissue cyst formation (stress-induced and spontaneous) in both species that also had a CRISPR phenotype score > −1, as we hypothesized that these factors would not be needed during tachyzoite growth. We also selected bradyzoite suppression factors (BSFs), genes whose transcript abundance decreased during tissue cyst formation, with low (≤ −1) CRISPR phenotype scores, as we predicted that these factors would be required for tachyzoite growth and therefore have poor fitness *in vitro*. After selecting candidates that fit these criteria, we had 16 final high priority candidates. (Figure 1C-D).

### Genetic ablation of of TGVEG_311100 impairs cyst formation *in vitro* while other candidate knockouts do not

To determine if our candidate genes were involved in tissue cyst formation, we disrupted each locus using CRISPR/Cas9. Specifically, we transfected parasites with a plasmid expressing Cas9-GFP under the control of the *T. gondii* SAG1 promoter as well as U6 gRNA expression cassette (Shen *et al*., 2017) that we modified to express a guide specific for our candidate gene or a non-targeting gRNA for control transfections. We co-transfected parasites with a linear insertion cassette encoding either the HXGPRT selectable marker for use in CEPΔHXGRPT-GFP-LUC (TgCEP) parasites or the DHFR-TS selectable marker. After cloning and validation using PCR, we successfully ablated TgME49_221840, TgVEG_207210, TgVEG_311100, and TgVEG_200385 (BFD1) (Figure S1).

We determined the impact of each gene disruption on alkaline-pH-stress-induced tissue cyst formation. After 48 hours of bradyzoite inducing conditions, we identified no significant differences in the number of DBA-positive vacuoles between the TgCEPΔ*221840* (Figure S2A) and TgVEGΔ*207210* (Figure S2B) knockouts relative to their passage-matched control. However, we did observe a significant decrease (*P*=0.0011) in the number of DBA-positive vacuoles for the TgVEGΔ*311100* parasites (∼25%) as compared to their WT controls (∼65%; Figure S2C). These data suggest that *TgVEG_311100* is an important gene for pH-induced cystogenesis and we named this factor Regulator of Cystogenesis 1 (ROCY1). Previous work identified BFD1 as necessary and sufficient for cyst formation in *T. gondii* (Waldman *et al*., 2020) and given its importance in differentiation we wanted to use this a means to calibrate ROCY1 cyst defects. Therefore, we generated TgVEG strains with a disruption in the BFD1 locus (Figure S2D) with the same passage history as our TgVEGD*ROCY1* parasite strain. We then performed a head-to-head comparison of tissue cyst formation in response to alkaline pH stress and found that both TgVEGΔ*BFD1* parasites failed to form DBA-positive cysts (Figure S2D) while TgVEGΔ*ROCY1* parasites formed significantly fewer DBA-positive cysts compared to the WT control (*P*=0.026). While the defect in DBA-positive vacuole formation was less robust in TgVEGΔROCY1 compared to TgVEGΔ*BFD1*, when we quantified DBA staining intensity we found a significant difference between TgVEGΔ*ROCY1* and WT parasites (P<0.001) and between TgVEGD*BFD1* and WT parasites (P<0.001; Figure S2E). Consistent with the intermediate phenotype of TgVEGΔROCY1, TgVEGΔBFD1 parasites also had significantly lower DBA staining intensity compared to TgVEGΔ*ROCY1* (P=0.006; Figure S2E-F). Together, these data suggest that ROCY1 plays an important role in the development of tissue cysts *in vitro*. Furthermore, the significantly higher intensity of DBA staining in the TgVEGΔ*ROCY1* parasites compared to TgVEGΔ*BFD1* parasites led us to hypothesize that ROCY1 may be in the same regulatory network as BFD1.

### *ROCY1* knockouts in a type II genetic background have reduced cyst formation *in vitro*

Given our dependence on oocyst production in cats to produce zoites of *H. hammondi* we used a well-characterized oocyst-forming parasite strain VEG for our comparative studies of *in vitro* development that led to the discovery of ROCY1. However, much of the *T. gondii* cyst formation studies to date have used strains belonging to the *T. gondii* type 2 haplotype like Prugniaud (PRU) (Soete *et al*., 1993) which also has been modified genetically in a variety of ways to permit more sophisticated genetic manipulations. To determine the impact of ROCY1 deletion in this genetic background we used the same reagents as in TgVEG to delete *ROCY1* and *BFD1* (again for calibration) in TgPru-GFP-LUC (Kim *et al*., 2007). We induced cyst formation using alkaline pH, low serum and atmospheric carbon dioxide and then stained for DBA 4 days later. We found that PRUΔ*ROCY1* had a robust cyst defect (only ∼10% of the observed vacuoles were DBA-positive cysts; Figure 2A-B), and as in TgVEGΔROCY1 had significantly lower DBA-staining intensity upon quantification. (Figure 2C). The defect in cyst formation was similarly robust in TgPRUΔBFD1, although no fully formed cysts were identified in TgPRUΔBFD1 (despite a lack of a statistically significant difference between them; Figure 2B). When DBA intensity was quantified in PruΔBFD1 it was also significantly lower than that in passage matched, wild type controls, but was statistically indistinguishable from PruΔROCY1 (*P*>0.05; Figure 2C). It is important to note here as well that even in control conditions (pH 7.2; left panel of Figure 2B-C) there was detectable DBA staining throughout the parasite vacuole in both wildtype and knockout parasite lines. Interestingly this staining in non-cyst forming conditions was significantly reduced in TgPRUΔROCY1 compared to WT controls (Figure 2C), and reduced, but not significantly so, in TgPRUΔBFD1 (Figure 2C). These data highlight the importance of ROCY1 and BFD1 in driving cyst formation during nutrient and pH stress, but also provide new insight into their role in driving spontaneous cyst formation (Jerome *et al*., 1998; Sokol *et al*., 2018).

**Figure 2:**
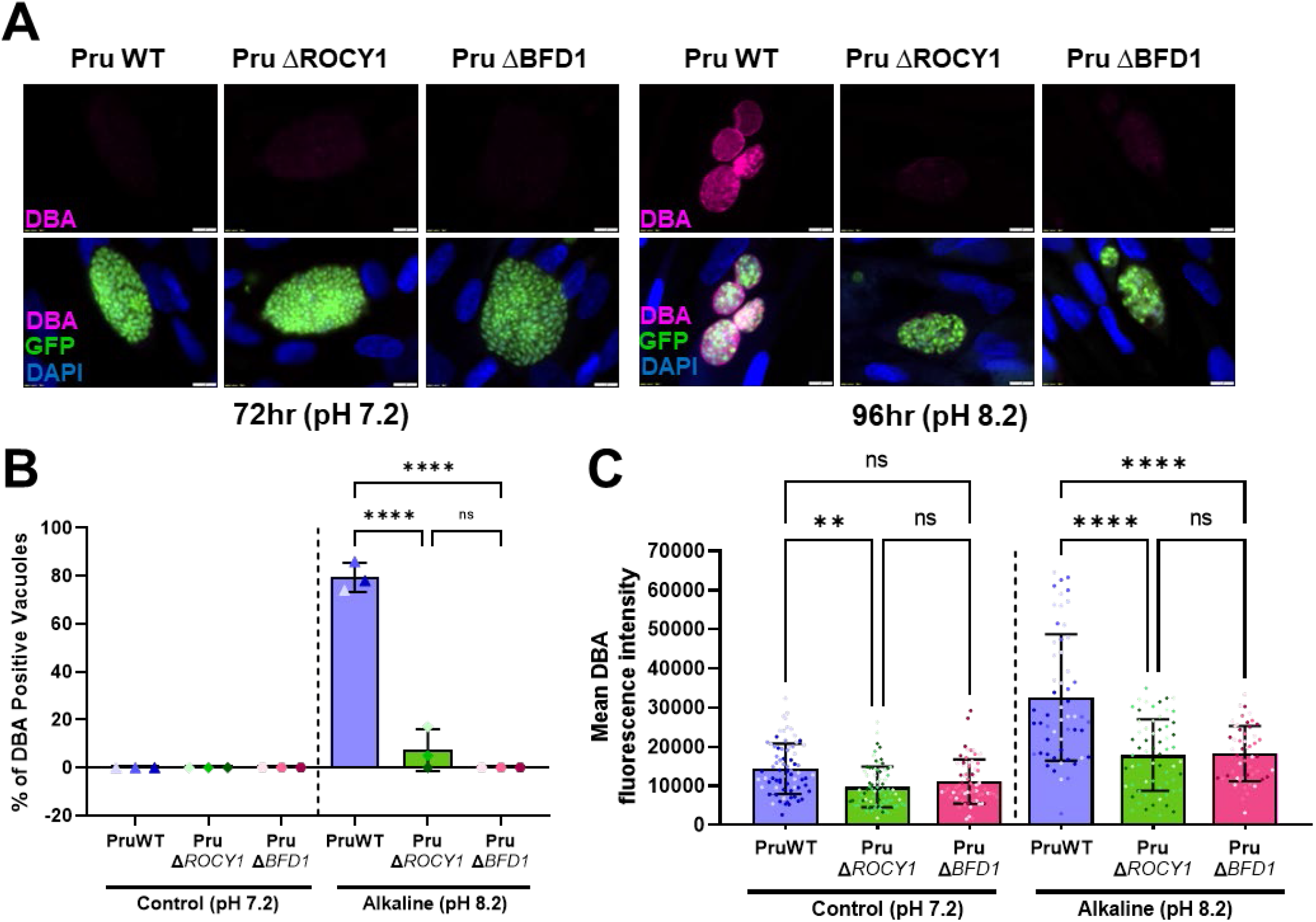
Genetic ablation of ROCY1 *T. gondii* PRU disrupts stress-induced cyst formation with similar efficacy as ablation of BFD1. A) Confluent monolayers of HFF cells in a 24-well plate were infected at an MOI of 0.3 with *T. gondii* (Pru WT, Δ*ROCY1* and Δ*BFD1*) that were each grown in pH 7.2 for 4 days and in pH 8.2 (bradyzoite switch induction conditions) for 3 days, fixed and stained with biotinylated DBA. B) Quantification of *T. gondii* infection described in A quantifying presence of DBA (+) cysts within counted vacuoles. Statistical significance was determined as in Figure S2 (****=P < 0.0001). Only Pru WT parasites formed significant numbers of cysts while ΔROCY1 and ΔBFD1 parasites were highly attenuated in stress-induced cyst formation. C) Detection and quantification of DBA fluorescent staining within parasites (mean intensity) as determined semi-automatically using KNIME software. Statistical significance was determined as in Figure S2 (****=P < 0.0001, **=P ≤ 0.01). Overall DBA intensity was significantly lower at high pH in both knockout strains and was significantly reduced in control pH conditions in PruΔROCY1 parasites.

### C-terminal tagging of ROCY1 inhibits cyst formation *in vitro*

As a first attempt to examine the localization of native ROCY1 we used CRISPR/CAS9 to tag the endogenous gene with a triple HA epitope in the TgPRUΔKu80ΔHXGPRT genetic background (Fox *et al*., 2011; Huynh and Carruthers, 2009). In PCR-and Sanger sequencing-validated tagged clones, we immunolocalized ROCY1 to punctate perinuclear structures in the parasite cytoplasm in both control (pH 7.2) and bradyzoite induction (pH 8.2) conditions (Figure 3A). Importantly staining intensity was qualitatively increased under high pH conditions (Figure 3A), consistent with the high transcript abundance of ROCY1 during pH-induced switching *in vitro*, *in vivo* bradyzoites (Pittman *et al*., 2014) and during spontaneous cyst formation in *H. hammondi* (Figure 1D). This staining patterns was similar to that observed for *T. gondii* ALBA proteins 1 and 2 (Gissot *et al*., 2013), a pair of RNA binding proteins in *T. gondii* involved in translational regulation. However when we quantified cyst formation (indicated by DBA staining surrounding the entire circumference of the vacuole) in two endogenously tagged clones we found that the C-terminal tag phenocopied the ROCY1 knockouts, switching at a frequency of <10% compared to over 70% in the wild type parent (P≤0.0001 for both clones; Figure 3B), suggesting that the C-terminal tag impaired ROCY1 function. To test this idea further we used CRISPR/CAS9 to add a single FLAG tag to the N-terminus of the *ROCY1* gene (again in TgPRUΔKu80ΔHXGPRT). Using immunofluorescence we observed that qualitatively the N-terminal FLAG tagged gene product localized identically to the C-terminally tagged gene product (Figure S3A), and that the FLAG-tagged product increased in overall abundance in bradyzoite induction conditions (Figure S3A-B). However in contrast to the C-terminal HA tagged strain, the N-terminally FLAG-tagged clones exhibited wild type levels of cyst formation under high pH conditions (Figure S3C). Taken together these data further confirm the importance of the ROCY1 gene in pH-induced cyst formation, and they also reveal a likely role for the C-terminus of ROCY1 in promoting pH-induced cyst formation.

**Figure 3:**
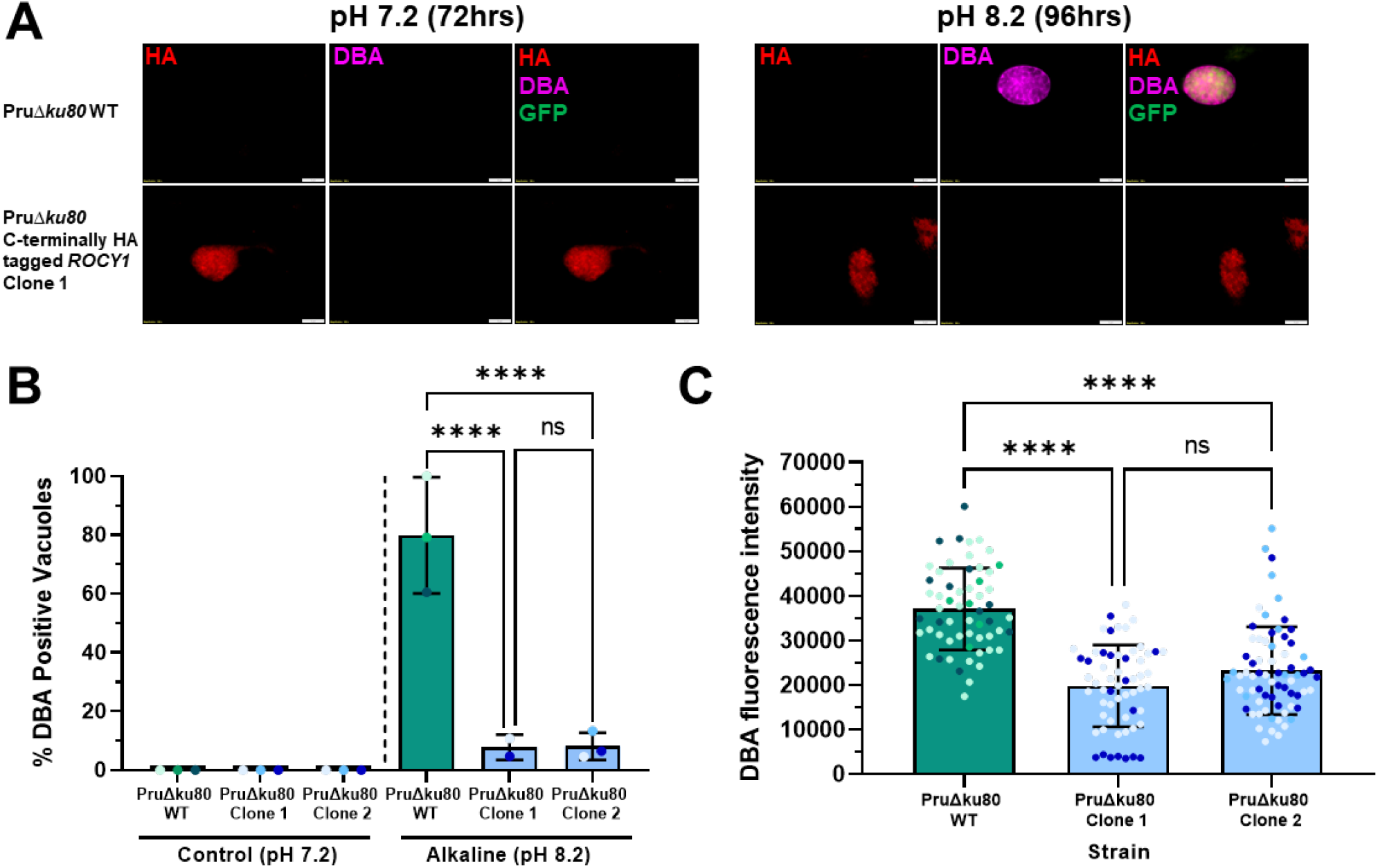
Endogenous C-terminal tagging ROCY1 in *T. gondii* PRUΔKu80 phenocopies the ROCY1 knockout with respect to stress-induced cyst formation. A) Confluent monolayers of HFF cells in a 24-well plate were infected at an MOI of 0.3 with *T. gondii* (PruΔ*ku80* WT and PruΔ*ku80* C-terminally HA tagged *ROCY1*) and grown in pH 7.2 for 4 days and in pH 8.2 for 3 days, fixed and stained with an anti-HA and biotinylated DBA. B) Quantification of the percentage of DBA-positive cysts from the infection described in A. Statistical significance was determined as in Figure S2. Stress-induced cyst formation is markedly impaired in clones with a 3X C-terminal HA tag at the ROCY1 locus. C) Detection and quantification of DBA fluorescence staining within parasites was found using KNIME software. Statistical significance was determined with a One-Way ANOVA test (****=P < 0.0001). Detection and quantification of DBA fluorescent staining within parasites (mean intensity) as determined semi-automatically using KNIME software. Statistical significance was determined as in Figure S2 (****=P < 0.0001, **=P ≤ 0.01). Overall DBA intensity was significantly lower in both 3X HA C-terminally tagged clones.

### ROCY1 is necessary and sufficient for both induced and spontaneous cyst formation

Based on our observation that N-terminal tags are tolerated by ROCY1 while C-terminal tags are not, we placed a single HA tag immediately upstream of the predicted start codon followed by a short linker sequence. Based on this information we made two complementation constructs with N-terminally-tagged ROCY1: one driving NHA-ROCY1 expression off of the putative ROCY1 promoter (Figure 4A; encompassing 2000 bp upstream of the start codon) and the other one driving NHA-ROCY1 using the GRA1 promoter (Figure 4B). The former was used to test if we could complement high pH-induced cyst formation in TgPRUΔROCY1 parasites, while the latter was to determine if ROCY1 was sufficient to induce cyst formation under tachyzoite growth conditions in TgPRUΔROCY1. We transfected TgPRUΔROCY1 parasites with each of these constructs as well as an empty vector containing only the GRA1 promoter and then grew them under MPA/Xanthine selection for 2-3 weeks prior to assessing the cyst formation capabilities of the populations under both tachyzoite (pH 7.2; Figure 4C) and bradyzoite (pH 8.2, 1% serum; Figure 4D) growth conditions. We then identified those parasites that were HA-positive and scored them as cysts or not based on DBA staining. Our data clearly show that TgPRUΔROCY1 parasites expressing NHA-ROCY1 off of the endogenous promoter could form cysts in response to high pH (Figure 4D; dark green bar in Figure 4E) at a significantly higher frequency than TgPRUΔROCY1 parasites transfected with the empty vector (Figure 4D; light green bar in Figure 4E). When we transfected parasites with GRA1-driven NHA-ROCY1, we found that even under tachyzoite conditions >30% of the HA-positive vacuoles had developed into DBA-positive cysts (Figure 4C; maroon bar in Figure 4E), and also that this “overexpression” ROCY1 construct was also capable of complementing the inducible cyst formation defects of PruΔROCY1 (Figure 4D-E). These data show that ROCY1 is not only necessary for inducible cyst formation *in vitro*, but also sufficient for cyst formation under control conditions when expressed off of a tachyzoite promoter.

**Figure 4.**
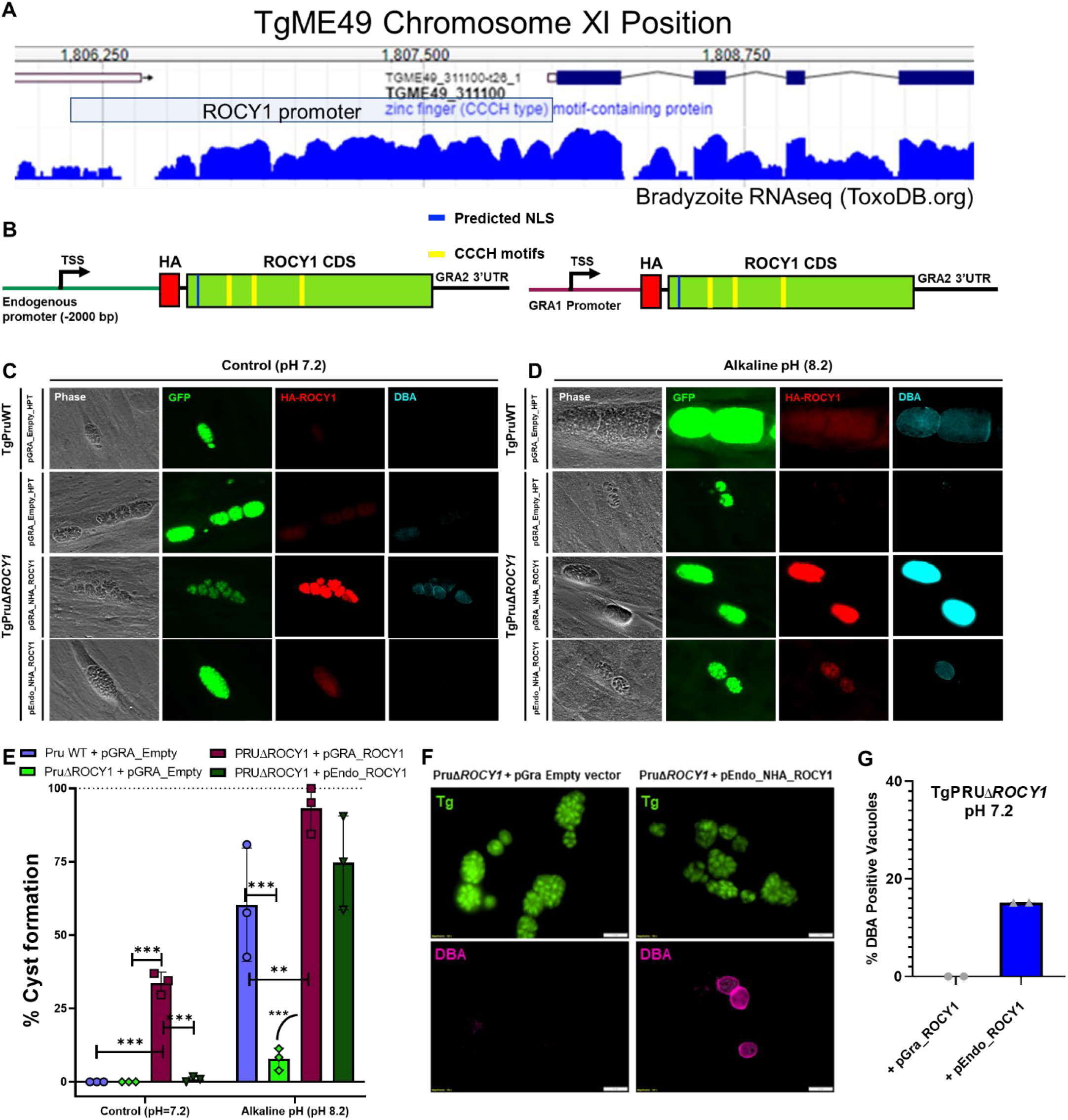
Complementation of ROCY1 restores tissue cyst formation phenotype. A) The ROCY1 locus as viewed on ToxoDB.org indicating the putative promoter used to complement ΔROCY1 parasites (light blue box) as well as publicly available RNAseq data representing ROCY1 transcript abundance and coverage. B) Schematic of complementation constructs using the endogenous promoter and a highly active GRA1 promoter. C-D) Representative images of TgPruA7 WT and TgPruA7Δ*ROCY1* parasites transfected with complementation constructs in C) control conditions and D) alkaline pH conditions E) Quantification of the average number of DBA positive cysts in 20 parasite containing fields of view in control and alkaline pH conditions. (N=3) Statistical significance was determined as in Figure S2. **P= 0.0025, ***P<0.0005, ****P<0.0001. Data show clear complementation of pH-induced cyst formation capacity in ΔROCY1 parasites with the pEndo_ROCY1 and pGRA_ROCY1 constructs (maroon and dark green bars), and the significant induction of cyst formation in control (pH 7.2) conditions in the presence of pGRA_ROCY1 (maroon bar on left.) F) Representative images of spontaneous cyst formation capacity of PruΔROCY1 parasites complemented with empty vector (left) or with the pEndo_ROCY1 construct (right). G) Quantification of “spontaneous” cyst formation in ΔROCY1 parasites by examining random fields of view and evaluating between 54 and 93 vacuoles across two biological replicates. While ΔROCY1 parasites transfected with the empty vector had no detectable DBA-positive cysts at pH 7.2 parasites complemented with the pEndo_ROCY1 construct formed cysts at a frequency of ∼15% (bar represents a total of 25 DBA-positive cysts out of 165 total vacuoles that were scored).

As we examined populations of TgPRUΔROCY1 and passage-matched TgPRU:WT parasites we observed that while wild type parasites formed DBA-positive cysts spontaneously at pH 7.2 at a rate of ∼1%, we rarely, if ever, saw DBA-positive cysts under control conditions in TgPRUΔROCY1, suggesting that “spontaneous” cyst formation is driven by ROCY1. To test this hypothesis we counted the number of DBA-positive cysts under normal growth conditions (pH 7.2) for TgPRUΔROCY1 and a clone of TgPRUΔROCY1 that was genetically complemented with pENDO_NHA_ROCY1 (see Figure 4B). In the TgPRUΔROCY1 parasite clone transfected with empty vector (pGRA_Empty), we did not identify a single DBA-positive cyst across 20 fields of view in each of 3 biological replicates (Figure 4F-G). In contrast, for TgPRUΔROCY1 complemented with the N-terminally tagged ROCY1 (pENDO ROCY1), we identified an average of 45+/-9 DBA-positive cysts (Figure 4G). This observation reinforces the dependence of cyst formation on ROCY1, in that parasites lacking this gene do not spontaneously enter the bradyzoite program (a common feature of many *T. gondii* strains; (Ferreira da Silva *et al*., 2008; Jerome *et al*., 1998)).

### At least one of three predicted CCCH-type zinc finger domains are required for ROCY1 induction of cyst formation

The *ROCY1* gene is predicted to encode a 920-amino acid protein that contains a predicted nuclear localization signal and 3 CCCH type Zinc finger domains (SMART accession SM000356; Figure 5A) and AlphaFold2 (Tunyasuvunakool *et al*., 2021) models of each of these motifs suggest that they functional given the orientation of the cysteine and histidine side chains capable of coordinating a single Zinc ion (Figure 5A). Based on this information we used multiple rounds of site-directed mutagenesis of the pGRA-NHA_ROCY1 construct to mutate 2 of the 3 cysteines to arginine and each histidine to lysine in all 3 CCCH motifs. After sequencing the entire CDS to verify the mutations and integrity of the ROCY1 coding sequence, we transfected TgPRUΔROCY1 parasites with the wild type pGRA:NHA_ROCY1 construct or the pGRA:NHA_ROCY1_CCCH mutant construct and grew the population under drug selection for 2 weeks. After growing the parasites on coverslips for 3 days we fixed, stained and then scored the HA+ vacuoles for their DBA-positive cyst phenotype under normal (i.e., pH 7.2) growth conditions. As expected, 63% of TgPRUΔROCY1 parasites expressing wild type NHA-ROCY1 were DBA-positive cysts, while not a single parasite vacuole (out of 105) expressing the CCCH mutant form of NHA-ROCY1 was scored as a DBA-positive cyst (Figure 5B). Representative examples of HA-positive vacuoles in both populations are shown in Figure 5C, and importantly show that the CCCH mutant form of NHA-ROCY1 has a similar localization pattern as the wild type form (Figure 5C). These data implicate the predicted CCCH motifs as being necessary for cyst formation in *T. gondii*, and based on the localization of tagged ROCY1 in cytoplasmic puncta as well as the propensity of proteins harboring CCCH-type zinc finger motifs to bind RNA, suggest that interactions with RNA by ROCY1 may be a key to its function in driving cyst formation.

**Figure 5:**
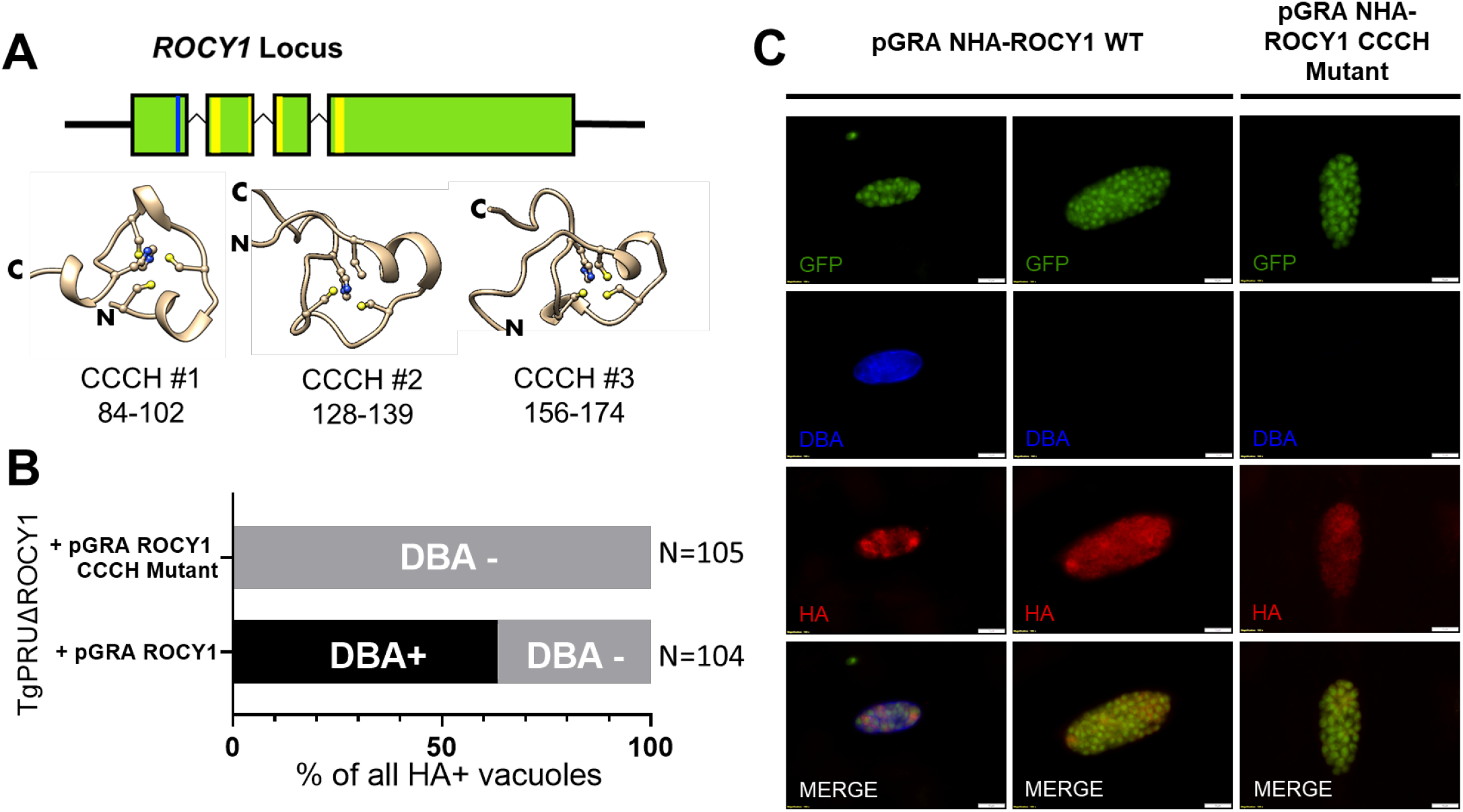
Mutating all 3 CCCH zinc finger binding domains of ROCY1 abolishes the effect of overexpressing ROCY1 in TgPRUΔROCY1 on cyst formation under control conditions. A) Schematic of the ROCY1 locus indicating the location of the putative NLS (blue bar) and CCCH motifs (yellow bars). Shown below are AlphaFold2 predictions for the 3 CCCH motifs, with each cysteine and histidine shown with side chains. N- and C-terminal directionality is indicated as are the locations of each CCCH motif (first Cysteine to the next histidine). B) Quantification of the percentage of DBA-positive cysts among HA-positive vacuoles expressing the WT pGRA NHA-ROCY1 (N=104) or the CCCH mutant form of pGRA NHA-ROCY1 (N=105) under control (pH 7.2) conditions. The CCCH mutations completely ablated the ability of pGRA NHA-ROCY1 to induce cyst formation. C) Representative images of DBA + and DBA - vacuoles containing PruΔROCY1 parasites expressing the wild type or mutant form of the pGRA_NHA-ROCY1 construct.

### Disruption of the *ROCY1* locus dramatically alters the alkaline pH-stress-induced transcriptional response

To better understand the role of ROCY1 in driving bradyzoite formation, we performed RNAseq on human foreskin fibroblasts (HFFs) infected with TgVEGΔ*ROCY1*, TgVEGΔ*BFD1*, and TgVEG WT parasites exposed to either control growth conditions (pH 7.2) alkaline pH stress conditions (pH 8.2) for 48 hours. Using principal component analysis (PCA) (Figure 6A), we found that separation along PC1 (62%) mostly represented the effect of pH and nutrient stress. All parasites grown in control conditions (WT, TgVEGΔ*ROCY1*, TgVEGΔ*BFD1*) clustered together along PC1 with minor separation in TgVEGΔ*ROCY1* and TgVEGΔ*BFD1* parasites from our TgVEG WT parasites along PCA2. Interestingly, we also saw a clear separation between TgVEG WT, TgVEGΔ*ROCY1*, and TgVEGΔ*BFD1* exposed to stressed conditions on both PC1 (62% of variance) and PC2 (9% of variance). Stressed TgVEG WT parasites were the most separated on PC1 compared to the control grouping but the least separated on PC2, while TgVEGΔ*BFD1* stressed parasites separated the least on PC1 but were the most separated on PC2 (Figure 6A). TgVEGΔ*ROCY1* parasites fell in the middle between the TgVEG WT and TgVEGΔ*BFD1* parasites exposed to stressed conditions. This transcriptional data is consistent with the slightly higher DBA-staining intensity that we observed in TgVEGΔROCY1 (Figure S2E-F).

**Figure 6.**
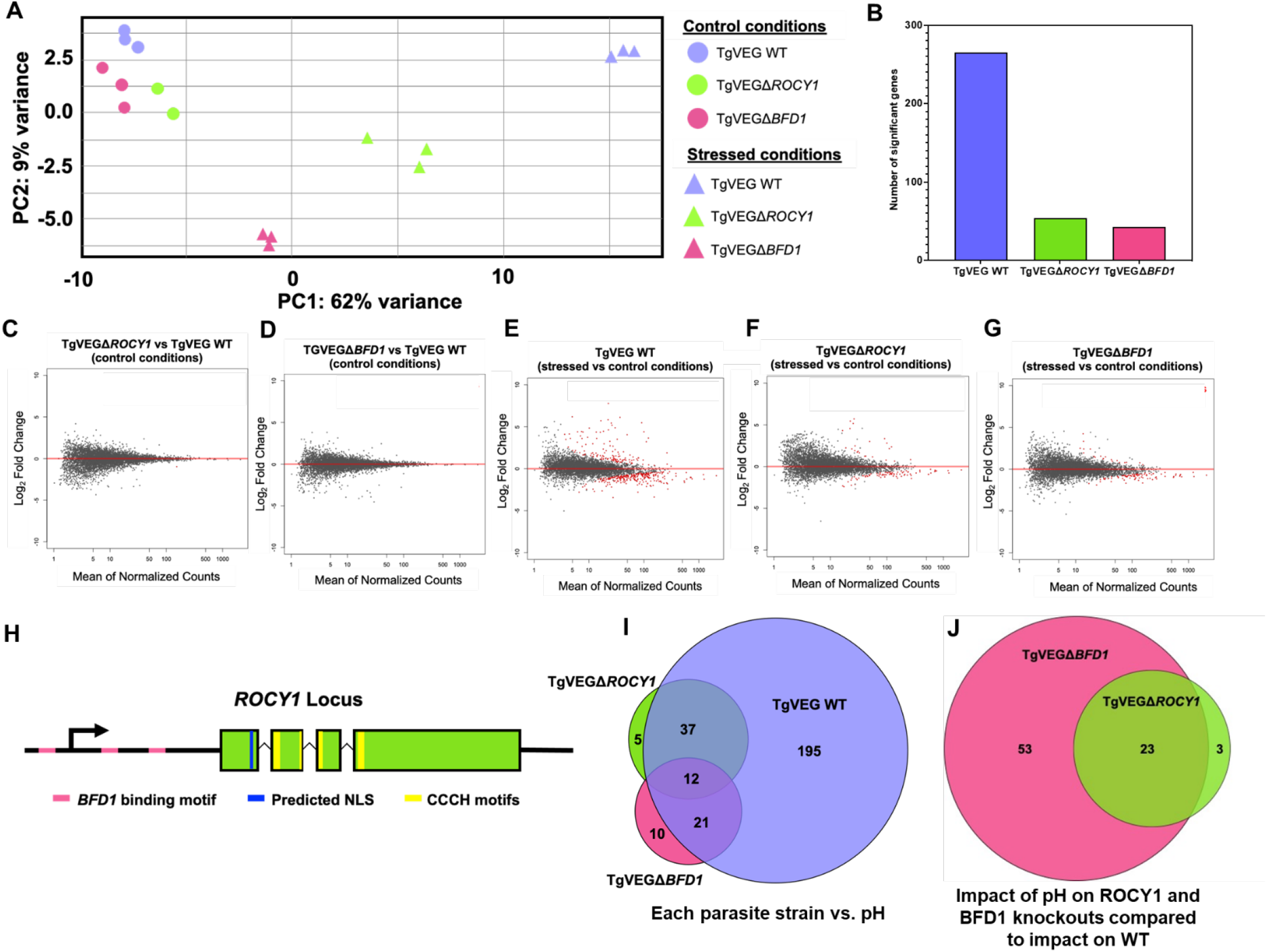
Disruption of the ROCY1 locus dramatically alters the alkaline pH-stress-induced transcriptional response. A) Principal components (PC) 1 and 2 of cells infected with TgVEG WT, TgVEGΔ*ROCY1,* and TgVEGΔ*BFD1* expose to either control (●) or alkaline pH stress (▲) conditions for 48 hours. (B) Number of significantly different transcript observed for infection with TgVEG WT, TgVEGΔ*ROCY1,* and TgVEGΔ*BFD1* (|Log_2_ Fold Change| ≥ 1, P_adj_ < 0.01). C-D) MA-plots representing changes in transcript abundance in TgVEGΔ*ROCY1* (C) or TgVEGΔ*BFD1* (D) infection compared to TgVEG WT infection in control conditions. Red dots (●) genes with a P_adj_ value < 0.01. E-G) MA-plots representing changes in transcript abundance in TgVEG WT (E), TgVEGΔ*ROCY1* (F), or TgVEGΔ*BFD1* (G) infections in alkaline stress conditions compared to control conditions. Red dots (●) genes with a P_adj_ value < 0.01. H) Diagram of ROCY1 locus showing predicted BFD1 binding sites both upstream and downstream of the predicted transcriptional start site, a predicted NLS, and 3 zinc finger CCCH motifs (second motif is split between exon 2 and 3). I) Venn diagram of genes with significant differences in transcript abundance in alkaline stress conditions compared to control conditions (|Log_2_ Fold Change| ≥ 1, P_adj_ < 0.01). J) Venn diagram of genes with significant differences in transcript abundance in TgVEGΔ*BFD1* (pink) and TgVEGΔ*ROCY1* (green) parasites compared to TgVEG WT parasites in alkaline stress conditions (|Log_2_ Fold Change| ≥ 1, P_adj_ < 0.01).

Based on differential expression analysis with DESeq2 we found that both TgVEGD*ROCY1* parasites and TgVEGD*BFD1* parasites exposed to control conditions had similar transcriptional profiles to TgVEG WT parasites as we did not identify any genes with significant changes in transcriptional abundance in either TgVEGΔ*ROCY1* parasites or TgVEGΔ*BFD1* parasites in comparison to WT (|Log_2_FC| > 1, P_adj_ < 0.01; Figure 6B-C). These data suggest that disruption of ROCY1 or BFD1 does not play a significant role during tachyzoite growth. We next looked at the transcriptional changes that occur in TgVEGD*ROCY1*, TgVEGD*BFD1*, and WT parasites in alkaline pH stress conditions (pH 8.2) relative to control conditions (pH 7.2). For WT parasites, as expected, we identified hundreds (265) of genes with significant differences in transcriptional abundance in response to treatment with alkaline stress compared to control conditions (|Log_2_FC| > 1, P_adj_ < 0.01) (Figure 3D-E). However, the alkaline-stress-induced transcriptional response was dramatically impaired for both the TgVEGD*ROCY1* parasites (54 genes with significant differences in abundance in stress conditions as compared to control conditions; Figure 3D,F) and TgVEGD*BFD1* parasites (43 genes; Figure 3D,H). These data suggest that BFD1 and ROCY1 regulate transcriptional abundance of the same set of genes or that BFD1 activates the expression of ROCY1 very early in the bradyzoite developmental pathway. When we examined the sequence upstream of the ROCY1 start codon (∼2,300 bp) we found 3 occurrences of a predicted BFD1 binding motif (CACTGG) within the previously identified CUT&RUN peaks (Waldman *et al*., 2020), two of which were found downstream of the predicted transcriptional start site (TSS) and another found upstream of the TSS (diagramed in pink in Figure 6H). The presence of BFD1 binding motifs in the putative ROCY1 promoter as well as our transcriptional data suggest that *ROCY1* transcription could be directly regulated by BFD1. To further investigate this hypothesis we quantified and compared ROCY1 and BFD1 transcript abundance in our RNAseq data and found that in stress conditions, *TgROCY1* transcript abundance is significantly lower (Log_2_FC = −1.46, P_adj_ < 0.001) in TgVEGD*BFD1* parasites relative to WT parasites while *TgBFD1* transcript is much more similar in abundance between stressed TgVEGΔ*ROCY1* and WT parasites (Log_2_FC = −0.58, P_adj_ = 0.04). Furthermore, we found that the majority of the dysregulated genes (23 of 26) in stressed TgVEGD*ROCY1* were also of different abundance in stressed TgVEGD*BFD1* (Figure 3J). Most of the dysregulated genes were of lower abundance (18 of 26 in TgVEGD*ROCY1* and 53 of 74 in TgVEGD*BFD1*) and many of those genes are known bradyzoite genes such as CST1, BPK1, and MCP4 that are components of the cyst wall (Buchholz *et al*., 2011; Tu *et al*., 2019; Zhang *et al*., 2001). Taken together these data suggest that ROCY1 functions as a positive regulator of bradyzoite development and is a likely downstream target of BFD1, and the significant overlap between BFD1- and ROCY1-dependent stress-induced transcripts suggests that these two factors are likely proximal in the bradyzoite development pathway.

### ROCY1 is necessary for brain tissue cyst formation during murine infections

Our *in vitro* data as well our transcriptional data suggest that ROCY1 is an important regulator of cystogenesis and is likely induced early in the cyst formation process. To determine if ROCY1 is also important for cyst formation *in vivo*, we infected CBA/J mice with TgVEG WT-GFP-LUC, TgVEGD*ROCY1*-GFP-LUC, and TgVEGD*BFD1*-GFP-LUC parasites in two separate experiments and monitored pathogenesis, morbidity and mortality using weight loss and *in vivo* bioluminescence imaging. In both experiments mice were first imaged at 3 h post-infection and we observed no significant differences in luciferase signal at this time point between strains. In the first experiment parasite burden was indistinguishable during the acute phase of infection (Figure 7A) across parasite strains, peaking at day 6 post-infection. We did not observe any detectable brain signal from parasites during establishment of the chronic phase of infection, a fact that we attribute to the hair and skin pigmentation in CBA/J mice and the lower brain parasite burden often associated with Type III strain infections. To examine brain parasite burden, we sacrificed mice at 4 weeks post-infection and used the brain to count GFP+/DBA+ cysts, to quantify the number of parasite genome equivalents per brain, to harvest RNA for RNAseq, and for immunohistochemical staining using anti-GFP antibodies. While mice infected with WT parasites harbored an average of 168+/-80 cysts/brain (Figure 7C), mice infected with TgVEGΔROCY1 had significantly fewer detectable brain cysts (2.8+/-5; P=0.0014), and cysts were detected in only 2 of the 8 mice that survived to 4 weeks post-infection. As expected based on previous studies on BFD1 (albeit in a type 2, rather than a type 3, genetic background; (Waldman *et al*., 2020)), TgVEGΔBFD1 were also deficient at forming cysts *in vivo*, and in these mice we failed to detect cysts in any of the mice (Figure 7C). When we quantified parasite genome equivalents using qPCR in a fraction of the brain, we found that in spite of the fact that *T. gondii* cysts were nearly undetectable in mice infected with TgVEGΔROCY1, there were no differences between any of the infections with respect to parasite genome equivalents in the brain (Figure 7D), and this was also true of TgVEGΔBFD1 (Figure 7D). We conducted this experiment a second time using similar infection and analysis regimens, but in this case we harvested mouse brains at 3 and 9 weeks post-infection. Parasite burden during the acute phase was again similar between strains, although we did detect significantly lower burden in TgVEGΔROCY1 parasites compared to wild type controls on days 7 and 8 post-infection (Figure S4B). In this experiment we did not detect any cysts at both 3 and 9 weeks post-infection in mice infected with TgVEGΔROCY1 or TgVEGΔBFD1 (Figure S4D,E), and we again detected significant amounts of parasite DNA in the brains of mice infected with all 3 parasite lines at 3 weeks post-infection (although we did detect significantly less DNA in brains of mice infected with TgVEGΔROCY1 parasites compared to controls (Figure S4E).

**Figure 7.**
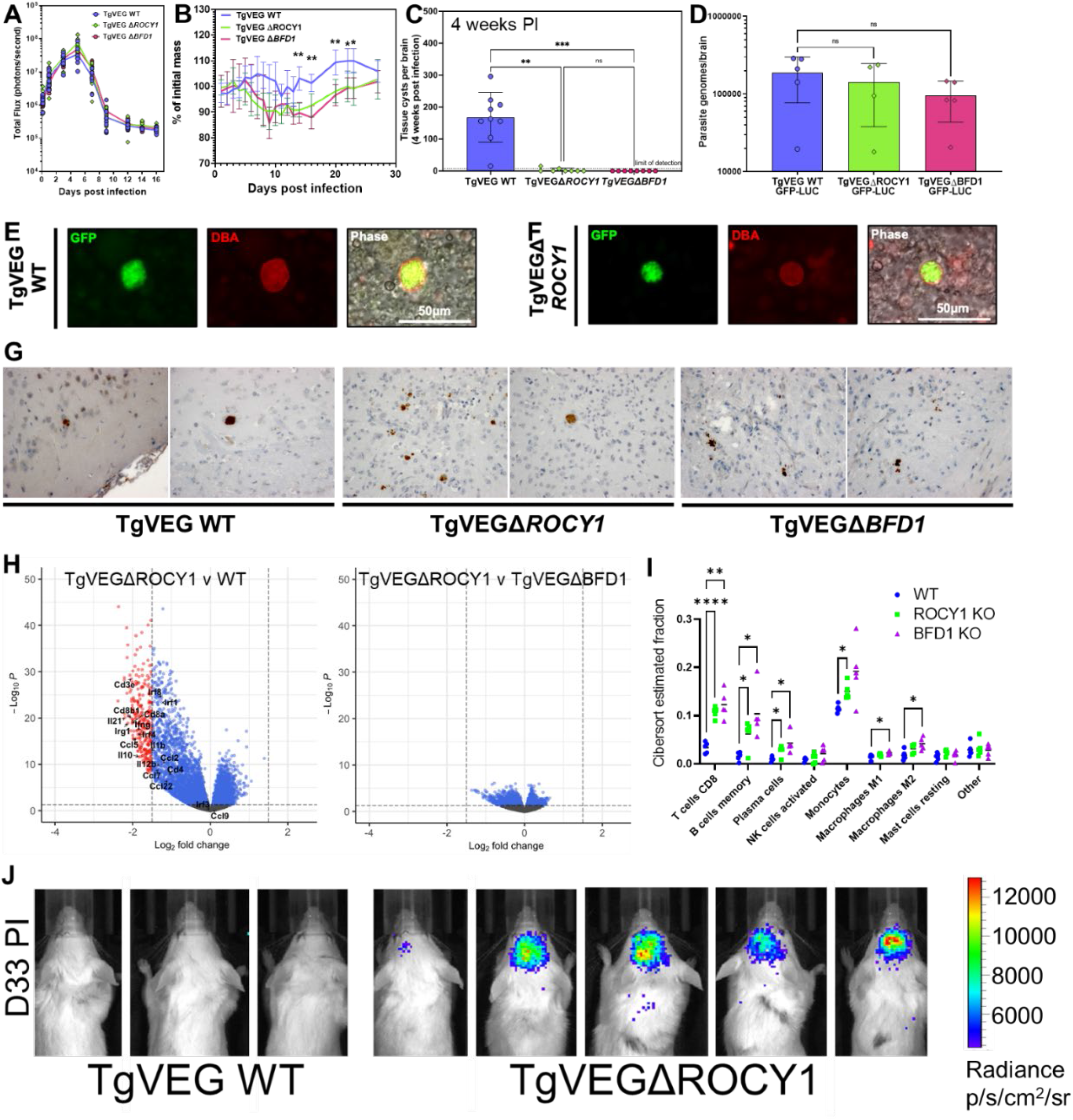
ROCY1 is necessary for tissue cyst formation and the establishment of chronic infection in mice. A) Quantification of bioluminescent imaging of mice infected with TgVEG WT-GFP-LUC, TgVEGΔ*ROCY1*-GFP-LUC, and TgVEGΔ*BFD1*-GFP-LUC parasite (N=6). Each point represents an individual mouse and the line represents the mean. B) Mouse weight loss over the course of the experiment in A, showing significantly higher weight loss in mice infected with ΔROCY1 or ΔBFD1 compared to wild type at multiple time points post-infection (asterisks indicate P<0.001 for both knockout strains). C) Brain tissue cysts burden observed in infected mice at 4 weeks post infection. Mean and standard deviation are plotted. Each point represents and individual mouse. Limit of detection = 7. Statistical significance was determined using a Kruskal-Wallis test with Dunn’s multiple comparisons. D) Quantification of parasite genomes per brain in infected mice at 4 weeks post infection. Mean and standard deviation are plotted. Each point represents an individual mouse. Statistical significance was determined using a one-way ANOVA with Tukey’s multiple comparison test on ΔC_t_ values (no statistically significant differences were observed). E-F) Representative images of TgVEG:WT (E) and TgVEGΔROCY1 (F) brain cysts stained with rhodamine-labeled DBA. G) Immunohistochemistry from intact brains using anti-GFP antibodies to localize T. gondii parasites. Two representative images from a single mouse for each infecting strain are shown. H) Volcano plots of sequenced RNA isolated from mouse brains 4 weeks post infection comparing transcripts between TgVEG WT versus TgVEGΔBFD1 infection, TgVEG WT versus TgVEGΔROCY1 infection, and TgVEGΔROCY1 versus TgVEGΔBFD1 infection. Log2 fold change cutoff = 1.5 (absolute value), p-value cutoff = 0.05 for significance. Transcripts labeled indicate a subset of immunity-related genes enriched during knockout infections as compared to TgVEG WT infections. I) Cibersort analysis of RNAseq data in (H) showing higher estimated proportions of CD8+ T cells as well as a other select cell types in mice infected with ΔROCY1 or ΔBFD1 parasites compared to controls. ***: P<0.001; **: P<0.01; *: P<0.05.

When we stained mouse brains from experiment 1 above using immunohistochemistry we found a qualitatively marked difference in vacuole structure between TgVEG:WT and TGVEGΔROCY1. While TgVEG:WT vacuoles were round in shape and contained within a single area (Figure 7G; left 2 panels), in TgVEGΔROCY1-infected brains we consistently saw vacuoles that were smaller in size and less compact in shape (Figure 7G; middle 2 panels). Again, this phenotype was recapitulated in TgVEGΔBFD1 infected mice (Figure 7G; right 2 panels). While vacuole phenotype was variable across sections and individual mice, this trend suggested to us that TgVEGΔROCY1 and TgVEGΔBFD1 parasites were persisting in the brain at 4 weeks post-infection in a state more reminiscent of tachyzoites than bradyzoites.

Since tachyzoites have a characteristic inflammatory signature that is distinct from slowly replicating bradyzoites, we reasoned that if ΔROCY1 parasites were indeed more similar to tachyzoites we may be able to detect this using sensitive measures of inflammation. Therefore we subjected subsamples of brains from 4-week infected mice (N=5 per parasite strain) to RNA sequencing and compared gene expression profiles using DESeq. Using PCA ΔROCY1 and ΔBFD1 brain RNAseq samples were both separated from wild type samples on the main principal component (Figure S5A) and were similarly adjacent based on unsupervised clustering (Figure S5B). Hundreds of transcripts were of different abundance in mouse brains infected chronically with TgVEGΔROCY1 compared to those infected with TgVEG:WT and most of these were shared between mice infected with both knockout parasite lines (Figure 7H and Figure S5C). The inflammatory signature of mice infected with ΔROCY1 and ΔBFD1 parasites was also significantly different as determined using gene set enrichment analysis (GSEA; (Subramanian *et al*., 2005)), where pathways like “HALLMARK INFLAMMATORY RESPONSE” and “HALLMARK IL2 STAT5” were similarly and significantly enriched based on a false-discovery rate of 0.05 (Figure S5D). We then used the informatics tool Cibersort (Newman *et al*., 2019) which can be used to infer percentages of different cell populations in bulk RNAseq data and that has recently been applied to brain RNAseq data from *T. gondii*-infected mice (Merritt *et al*., 2020). Given our interest in the inflammatory profile of mice infected with wild type and cyst-deficient *T. gondii* (ΔROCY1 and ΔBFD1), we used the precomputed “LM22” as the signature gene file and used it to estimate the relative proportions of 22 immune cell types using our brain RNAseq data. Of the 22 cell types queried CD8+ T cells were predicted to be significantly higher in abundance in both ΔROCY-infected (P<0.0001) and ΔBFD1-infected (P<0.01) mice based on brain RNAseq data (Figure 7I). These data are consistent with the idea that by 4 weeks post infection wild type TgVEG parasites are causing significantly less inflammation compared to ΔROCY1 and ΔBFD1 parasites, and that there may be more CD8+ T cells in the brains of mice infected with the mutant parasites to combat was is being sensed as an acute, rather than chronic, infection. These data are also consistent with mouse imaging studies from BALB/c mice (which permit more sensitive detection of brain-associated parasites). We infected mice with either TgVEG:WT or TgVEGΔROCY1 parasites and imaged their heads dorsally during the chronic phase of infection. We observed a consistently higher luminescent signal in the brain of TgVEGΔROCY1-infected mice starting on day 13 post-infection, and found that detectable luminescence persisted in TgVEGΔROCY1-infected mice for up to 33 days (Figure 7J). This was in stark contrast to wild type *T. gondii* VEG, which we failed to detect in the head area at 33 days PI (Figure 7J) and at any time point after day 18 post-infection (Figure 7J). Taken together these data illustrate a distinct outcome for the transition to the chronic phase of infection in the ΔROCY1 and ΔBFD1 parasite lines and this is likely due to their failures to successfully form cysts.

### *Toxoplasma* parasites that fail to form cysts are still capable of reactivation for up to 5 months post-infection

Even though we failed to observe tissue cysts in the brains of either the TgVEGD*ROCY1*-GFP-LUC and TgVEGD*BFD1*-GFP-LUC parasites, we did observe parasite DNA and signatures of an inflammatory response. A long-standing hypothesis is that cyst formation itself is required for long-term persistence of parasites in animal tissues, and therefore if one could somehow inhibit cyst formation then the immune response would rapidly clear these parasites that lack the protection afforded by the cyst wall and “quiescent” nature of the bradyzoite containing cyst. We reasoned that we could test this largely accepted concept by performing reactivation experiments in mice infected with our luciferase-expressing TgVEGΔROCY1 and ΔBFD1 parasites. To do this we infected mice with TgVEG WT, TgVEGD*ROCY1,* and TgVEGD*BFD1* parasites and assigned them to either the control group (N=6) or the reactivation group (N=6). On day 30 post infection, mice in the reactivation group were given dexamethasone in their drinking water to induce reactivation and subsequent reactivation was monitored using bioluminescence imaging (Nicoll *et al*., 1997; Saeij *et al*., 2005). We monitored reactivation by imaging the infected mice on multiple days after dexamethasone treatment (Figure 9A-B). As expected based on prior work we observed a significant increase in luciferase signal in dexamethasone-treated mice infected with TgVEG WT parasites as compared to non-treated control mice at day 4 post-dexamethasone treatment (P_adj_=0.038) and at day 8 post dexamethasone treatment (P_adj_=0.047) indicative of reactivation. We also observed clear evidence for reactivation in mice infected with TgVEGΔROCY1 (on day 8 post-dexamethasone treatment) and TgVEGΔBFD1 (on days 6-14 post-treatment; Figure 9A-B). We also looked at the difference in the survival of mice infected with TgVEG WT, TgVEGΔ*ROCY1,* and TgVEGΔ*BFD1* and treated with dexamethasone. We found that dexamethasone treated mice infected with TgVEGΔROCY1 and TgVEGΔ*BFD1* died significantly sooner than dexamethasone treated mice infected with TgVEG WT (P=0.014 and 0.002, respectively; Figure S6A). We repeated this experiment a second time and obtained similar results. Again *T. gondii* parasites reactivated regardless of parasite strain (Figure 9C), and again mice infected with either TgVEGΔROCY1 or TgVEGΔBFD1 succumbed to the infection significantly sooner (P<0.0001) compared to those infected with wild type parasites and treated with dexamethasone (Figure 9C and S6B). Finally we also tested the reactivation kinetics in mice 5 months post-infection to see if *T. gondii* ΔROCY1 and ΔBFD1 parasites were capable of reactivating long-term chronic infection. Again, as for parasites infected for 30 days post-infection we found that all 3 strains were capable of reactivating at this late time point (Figure 9D-E) within a week of the start of dexamethasone treatment. These data clearly show that *T. gondii* parasites with dramatic defects in cyst formation are paradoxically capable of long-term persistence in a mouse model as revealed by their ability to reactivate and disseminate after immune suppression.

**Figure 9.**
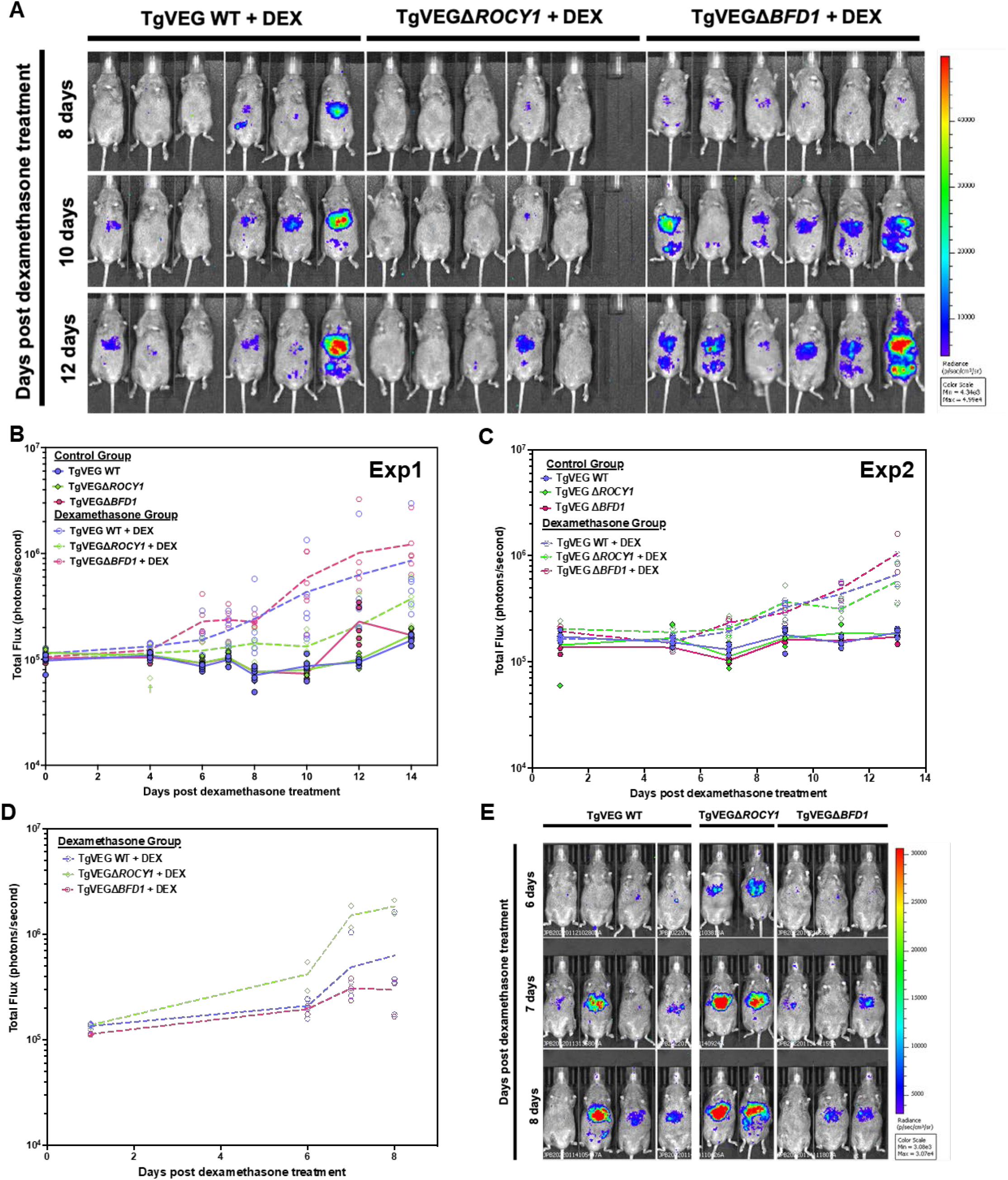
TgVEGΔ*ROCY1* and TgVEGΔ*BFD1* parasites can be reactivated following chronic infection. A) Select bioluminescent images showing parasite-derived luciferase signal in CBA/J mice infected with TgVEG WT, TgVEGΔ*ROCY1,* and TgVEGΔ*BFD1* parasites and treated with dexamethasone (20mg/L) provided in their drinking water beginning at 30dpi. B) Quantification of bioluminescent imaging of mice infected with TgVEG WT, TgVEGΔ*ROCY1,* and TgVEGΔ*BFD1* parasite belonging to either the control group (N=6 per parasite strain) or the dexamethasone treatment group (N=5 −6 per parasite strain). Each point represent an individual mouse and the line represents the mean. The double cross symbol (☨) represents death due to handling/anesthesia. Dashed line annotations represent dexamethasone group compared to control group for each parasites strain. Blue annotations represent TgVEG WT-GFP-LUC, green annotations represent TgVEGΔ*ROCY1*-GFP-LUC, and pink annotations represent TgVEGΔ*BFD1*-GFP-LUC. B-C) Quantification of bioluminescent imaging of mice infected with TgVEG WT, TgVEGΔ*ROCY1,* and TgVEGΔ*BFD1* parasite belonging to either the control group (B, Exp1: N=6, C, Exp2: N= 3-5 per parasite strain) or the dexamethasone treatment group (B, Exp1: N= 5-6, C, Exp2: N=3 per parasite strain) reactivate at 30dpi for B and 28dpi for C. Each point represent an individual mouse and the line represents the mean. Dashed line annotations represent dexamethasone group compared to control group, solid line, for each parasites strain. Blue annotations represent TgVEG WT-GFP-LUC, green annotations represent TgVEGΔ*ROCY1*-GFP-LUC, and pink annotations represent TgVEGΔ*BFD1*-GFP-LUC. (See stat’s table on next slide, EXP1 had significant reactivation for all Exp2 didn’t for ROCY1) D) Quantification of bioluminescent imaging of mice infected with TgVEG WT, TgVEGΔ*ROCY1,* and TgVEGΔ*BFD1* parasites following dexamethasone treatment (20mg/L provided in drinking water) after 5 months of infection. E) Select bioluminescent images showing parasite-derived luciferase signal at days 6, 7, and 8 post dexamethasone treatment (day 153, 154, and 155 post infection).

## Discussion

The formation of bradyzoites is necessary for the establishment of chronic infection which is fundamentally important for persistence of the parasite within its host and transmission to the next host (Dubey, 1998, 2010; Jeffers *et al*., 2018; Tenter *et al*., 2000). Despite the importance of bradyzoites, our understanding of the developmental pathways and the regulatory networks used by *T. gondii* to differentiate into these life stages is only starting to be elucidated. By using cross-species comparisons with *H. hammondi* at critical developmental time points, we have identified a previously uncharacterized parasite gene, *ROCY1*, that is important for tissue cyst formation *in vitro* and *in vivo* and is both necessary and sufficient for cyst formation.

Given the importance of the tissue cysts for allowing parasites to reside in host tissue without being continually subjected to immune pressures during chronic infection (Dubey *et al*., 1998; Weiss and Kim, 2000), we were surprised that parasites lacking ROCY1 (and BFD1, for that matter) were as capable of reactivation and to cause systemic and lethal disease following immune suppression with dexamethasone for up to 5 months post-infection. To our knowledge, no other reactivation studies have been performed with parasites that are incapable of forming cysts, and this observation calls into question the importance of cyst formation itself in parasite persistence. Our data clearly show that TgVEGΔROCY1 parasites were as capable at colonizing and persisting in the brain as their wild type counterparts. Moreover, based on the fact that we detected very few cysts in our brain preparations suggests that any parasites present in the brain were NOT cysts, as they were disrupted by the relatively mild process of straining brain material through a 100 μm cell strainer (a standard way we and others purify cysts; (English and Boyle, 2018)). However, host immunity is thought to kill the majority of tachyzoites during the acute phases of infection and is hypothesized to promote the survival of bradyzoites by either eliminating tachyzoites or by inducing bradyzoite development (Skariah *et al*., 2010). If the majority of tachyzoites are killed by host immunity, the ability of TgVEGΔ*ROCY1* and TgVEGΔ*BFD1* to persist and then reactivate when treated with dexamethasone deep into the chronic phase of infection suggests that 1) some tissue cyst formation is occurring in the infected mice at levels below the detection threshold in the brain, 2) tissue cysts are forming in other tissues (such as muscle, spleen, or liver) instead of forming in the brain, or 3) these parasites can survive in the mouse despite being unable to form tissue cysts. Our data point most frequently at explanation #3, which means that that brain-and other tissue-resident ΔROCY1 and ΔBFD1 parasites are more like tachyzoites than anything else. This idea finds further support in our host RNAseq and Cibersort data showing significantly higher levels of inflammation in mice chronically infected with ΔROCY1 or ΔBFD1 parasites, and the increased speed with which mice succumbed to reactivated infection when previously infected with knockout parasites compared to wild type controls. It is possible, therefore, that cyst formation (or even the transition from the tachyzoite to the bradyzoite irrespective of the formation of a cyst wall) is NOT required for *T. gondii* parasites to survive for long periods of time in the infected host.

Our *in vitro* cyst induction assays along with our transcriptional analysis suggest that ROCY1 is critical for bradyzoite development. Our transcriptional data suggests that ROCY1 has a role in increasing the transcriptional abundance of key bradyzoites transcripts, such as DnAK-TPR, CST1, BPK1, and MCP4. Some of these bradyzoite specific transcripts have also been shown to be regulated by other known transcription factors such as the bradyzoite transcriptional repressors AP2IX-4 (Huang *et al*., 2017) AP2IV-4 (Radke *et al*., 2018), AP2IX-9 (Radke *et al*., 2013), and bradyzoite transcriptional activators AP2XI-4 (Walker *et al*., 2013) and AP2IV-3 (Hong *et al*., 2017). We identified extensive overlap in the dysregulated genes in TgVEGΔ*ROCY1* parasites as compared to TgVEGΔ*BFD1* parasites, implying that ROCY1 functions in the same pathway as BFD1. ROCY1 is at least partially regulated by BFD1, a premise also supported by the presence of multiple BFD1 binding motifs in the promoter of ROCY1 (Waldman *et al*., 2020). However observations in our present study suggest that ROCY1 is not strictly a downstream effector wielded by BFD1. Most importantly, we observed that overexpressing ROCY1 (i.e., by using a “tachyzoite” promoter like that of GRA1) is sufficient to induce cyst formation (Figure 4C-E). Second, we observed that spontaneous cyst formation (i.e., cyst formation under control or “pH 7.2” conditions) depends on ROCY1 (Figure 4F-G). This suggests some interdependence of each of these factors on the other, a suggestion that finds important support in a co-submitted study from the Lourido lab (Licon *et al*. “A positive feedback loop controls *Toxoplasma* chronic differentiation”) where they found that ROCY1 binds directly to the BFD1 transcript in a stress-dependent fashion. Overall the findings from our research group as well as that of Licon *et al*. have demonstrated at a minimum that ROCY1 is very proximal to BFD1 in the bradyzoite developmental pathway, and provide extensive evidence that their transcripts and/or gene products may interact genetically and/or physically.

The predicted amino acid sequence of ROCY1 indicates that this protein contains 3 tandem CCCH zinc finger domains and these are absolutely essential for the ability of this gene to induce cyst formation under overexpression conditions. While zinc finger proteins are typically viewed as DNA-binding transcription factors, proteins containing CCCH zinc finger domains are known to bind RNA (Fu and Blackshear, 2017; Hall, 2005). Interestingly, RNA binding proteins with CCCH zinc finger domains have been implicated in the regulation of growth and stress responses in plants, response to oxidative stress in mammalian cells, and the regulation of immune responses (Fu and Blackshear, 2017). CCCH zinc fingers have also been known to promote RNA deadenylation and degradation by binding to AU rich elements in the 3’ UTRs of mRNA transcripts (Lai *et al*., 2000). *T. gondii* RNA binding proteins, such as TgAlba1, and TgAlba2, are involved in bradyzoite development and function in translational control (Gissot *et al*., 2013), although it should be noted that the TgΔAlba1 cyst formation phenotype in mice was not as dramatic as that observed here for TgVEGΔROCY1. These findings, along with the critical requirement for at least one of the 3 CCCH zinc fingers based on our site-directed mutagenesis studies, suggest an intriguing role for ROCY1 in regulating transcripts of key developmental genes induced during the alkaline pH stress response in *T. gondii*. A critical next step will be to determine both the subtype (e.g.., mRNA, small RNA, non-coding RNA, ribosomal RNA) and identity of the RNA molecules that are specific substrates for ROCY1.

Taken together, these findings demonstrate that ROCY1 is an important parasite factor for bradyzoite development in *T. gondii* and contributes to the development of a gene regulatory network used by these parasites to initiate the changes required for effective stage conversion. Future studies investigating the role of these factors in bradyzoite development in other parasite species, such as *H. hammondi*, that also rely on bradyzoite development for success, will help compare the strategies used by each species to progress to the bradyzoite life stage. Understanding the complexities of bradyzoite development associated between species has the potential to shed light on the fundamental differences attributed to each parasite’s unique life cycle.

## Materials and methods

### Host cell and parasite strains

*Toxoplasma gondii* strain VEG (TgVEG) (Dubey, 2001) and *Hammondi hammondi* strain HhCatAmer (HhAmer) (Frenkel and Dubey, 1975) were isolated from oocysts from experimentally infected cats provided by J.P Dubey as previously described (Sokol *et al*., 2018, 2020). The *Toxoplasma gondii* strain CEPΔHXGRPT-GFP-LUC (TgCEP) (Kamau *et al*., 2011) was also used. All *T. gondii* parasites were maintained through passage in Human foreskin fibroblasts (HFFs) cultivated with DMEM supplemented with 100U/mL penicillin, 100µg/mL streptomycin, 2mM L-glutamine, and 10% FBS (cDMEM) and maintained at 37 degrees C, 5% CO2.

### Oocyst excystation

Sporozoites were harvested from oocysts as previously described (Sokol *et al*., 2018, 2020). Following excystation, sporozoites were used to infected confluent monolayers of HFFs grown in T-25 tissue culture flasks and incubated at 37 degrees C, 5% CO2 for 24 hours. Following the initial 24-hour incubation, the infected monolayers were scrape, syringe lysed with a 25G needle, and filtered through a 5µm syringe driven filter. Zoites were then pelleted and counted to set up subsequent infections.

### RNAseq of *H. hammondi* (HhAmer) and *T. gondii* (TgVEG) in control and bradyzoite induction conditions

HFFs were infected with 24 hour “zoites” at an MOI of 0.5 for *T. gondii* and 8.5 for *H. hammondi*. After 48 hours (3DPE), infected host cells were exposed to either bradyzoite induction media, pH 8.2, (Naguleswaran *et al*., 2010) or control media, cDMEM pH 7.2, for 48 hours. At 48 hours, the infected host cells were washed with PBS and RNA was harvested using the RNeasy Mini Kit 9(Qiagen). Total isolated RNA was processed for next generation sequencing as previously described (Sokol *et al*., 2018). CLC Genomics Workbench was used to map all fastq reads to either the *T. gondii* genome or the *H. hammondi* genome as previously described (Sokol *et al*., 2018). Total gene counts were exported from CLC and filtered so that only genes with a total of 10 reads across all parasite-specific samples were used for differential expression analysis using DESeq2 (Varet *et al*., 2016). A total of 7,182/8,920 genes were analyzed for *T. gondii* and 6106/7266 for *H. hammondi*. Venn diagrams were created with BioVenn (Hulsen *et al*., 2008) and Venn Diagram Plotter software (Pacific Northwest National Laboratory). Heat maps were generated with MeV 4.9.0 (Saeed *et al*., 2003).

### Disruption of candidate gene loci

The CRISPR/Cas9 gene editing strategy was used to disrupt loci of candidate genes. For each candidate gene specific guide RNAs (gRNAs) were designed using the E-CRISP design tool (*Toxoplasma gondii* genome, medium setting). The gene specific gRNAs were incorporated into a version of the the pSAG1::CAS9-U6::sgUPRT plasmid provided by the Sibley lab (Shen *et al*., 2014) that was engineered to so that the UPRT gRNA was replaced with an PseI and FseI restriction site (Rudzki *et al*., 2020) (pCRISPR_ENZ) using Q5 mutagenesis (NEB) and verified with Sanger sequencing. Linear repair constructs encoding either the HXGPRT and DHFR-TS selectable markers under the control of the DHFR 5’ and 3’ UTR were amplified from either the pGRA_HA_HPT (Saeij *et al*., 2006) or pLIC-3HA-DHFR (Sugi *et al*., 2016) using Platinum® *Taq* DNA Polymerase High Fidelity with High Fidelity PCR buffer and 110ng of template per reaction according to the manufacturer’s reaction conditions. Approximately 10×10^6^ parasites were transfected with 25µg of the NheI-HF linearized CRISPR plasmid and 2.5µg of the appropriate linear repair cassette in 800µL of Cytomix (10mM KPO_4_, 120mM KCL, 5mM MgCl_2_, 25mM HEPES, 2mM EDTA) supplemented with 2mM ATP and 5mM glutathione using a BTX electroporation system – Electro Cell Manipulator 600 (2.4kV Set Charging Voltage, R3 Resistance timing). Transfection reactions were incubated at room temperature for 5 minutes before infecting confluent monolayers of HFFs. After ∼24 hours, the infected host cells were grown in selection media containing cDMEM supplemented with mycophenolic acid (25µg/mL) and xanthine (50µg/mL) for TgCEP or cDMEM with 1µM pyrimethamine for TgVEG to select for parasites that incorporated the repair cassettes containing the selectable markers. After a stable population of parasites capable of growing in selection media was obtained, clonal population were obtained through serial dilution. Genomic DNA was isolated from the clonal populations and used to verify disruption of the candidate gene locus with the selectable marker with PCR (2x Bio^TM^Mix Red used according to manufacturer’s specifications, 25-50ng of genomic DNA per reaction) Amplified products were visualized with 1% agarose gels run at 90 volts for 25-40 minutes. The GeneRuler 1kb Plus DNA Ladder was used to estimate the size of the amplified products.

### Alkaline pH-stress-induced tissue cyst formation assays

Confluent monolayers of HFFs grown on acid-etched coverslips in 24-well plates were infected with *T. gondii* at an MOI ranging from 0.25 to 1 depending on experiment. MOIs were the same for all tissue cyst formation assays performed on the same day. Parasites were grown for 48 hours and then their media was replaced with either control media (cDMEM pH 7.2) or bradyzoite induction media, pH 8.2, (Naguleswaran *et al*., 2010). Parasites grown in control conditions were incubated at 37 degrees C, 5% CO_2_ and parasites grown in alkaline pH stress conditions were incubate at 37 degrees C with ambient CO_2_ (0.03%). Control or bradyzoite induction media was replaced every 24 hours. After different times of alkaline pH stress, infected coverslips were washed 2X with PBS, fixed with 4% paraformaldehyde in PBS, washed 2x with PBS, and stored in PBS at 4 degrees C until immunostaining was performed.

### Immunofluorescence assays

Fixed coverslips were blocked with 5% BSA in PBS with 0.15% Triton-X 100 for 1 hour at room temperature. Coverslips infected with TgVEG WT, TgVEGΔ311100 (ROCY1), TgVEGΔ200385 (BFD1), TgVEGΔ207210, and TgCEPΔ221840 were stained with a primary polyclonal anti-MAF1b from mouse serum (1:1,000 dilution). After primary staining, the coverslips washed with PBS, stained with secondary goat anti-mouse Alexa-fluor 488 antibody and Rhodamine-labeled or biotinylated *Dolichos Biflourus* Agglutin (1:250) and washed with PBS. For biotinylated DBA this was followed up by incubating in streptavidin coupled to either AlexaFluor 405 or AlexaFluor 647. Coverslips were then mounted using ProLong Diamond Antifade mountant or Vectashield (both with and without DAPI) and allowed to cure overnight at room temperature if necessary. Coverslips were blindly observed with a 100X objective on an Olympus IX83 microscope with cellSens software. The percentage of DBA-positive vacuoles was generally determined by counting vacuoles from 15 randomly chosen, parasite containing fields of view, although in some cases parasites were first identified to be expressing a transgene of interest prior to scoring.. Images were exported as .tiff files and average fluorescence intensity per pixel was quantified using ImageJ software and corrected for background fluorescence intensity.

### RNAseq of TgVEGΔ*ROCY1* and TgVEGΔ*BFD1* in control and bradyzoite induction conditions

Confluent monolayers of HFFs grown in 24-well plates were infected with TgVEG WT, TgVEGD*ROCY1,* and TgVEGD*BFD1* parasites at an MOI of 0.5. Parasites were grown in control conditions (cDMEM pH 7.2, 37 degrees C, 5% CO_2_) for 48 hours. After 48 hours the media was replaced with either control media (cDMEM pH 7.2) or bradyzoite induction media, pH 8.2, (Naguleswaran *et al*., 2010). Infected host cells grown in control conditions were incubated at 37 degrees C, 5% CO_2_ and infected host cells grown in bradyzoite induction media were incubated at 37 degrees C with ambient CO_2_ (0.03%). Control or bradyzoite induction media was replaced every 24 hours. After 48 hours of growth in bradyzoite induction conditions, all infected host cells were washed with PBS and total RNA was harvested using the RNeasy Mini Kit 9 (Qiagen). This RNA was then processed for next generation sequencing as previously described (Sokol *et al*., 2018). Fastq files were first mapped to the human genome to remove host transcripts, Hg38 using CLC Genomics Workbench (default mapping settings for reverse strand sequencing, similarity fraction adjusted to 0.95) and a file of unmapped reads was generated for each sample. Unmapped reads were then mapped to the *T. gondii* genome (TgME49 v38) (default mapping settings for reverse stand sequencing). Total gene counts were exported from CLC and filtered so that a gene was only included for differential expression analysis using DESeq2 (Varet *et al*., 2016) if there were a total of 30 or more reads across all samples. A total of 5,970/8,920 genes were analyzed. Venn diagrams were created with BioVenn (Hulsen *et al*., 2008) and Venn Diagram Plotter software (Pacific Northwest National Laboratory).

### Endogenous tagging

C-terminal endogenous tagging of loci was performed in TgPRUΔKu80ΔHXGPRT by targeting the 3’UTR of ROCY1 with a specific gRNA cloned into the pUniversal-CAS9 plasmid (Addgene #52694) and then co-transfected with a homology repair cassette amplified from the pLIC-HXGPRT plasmid (as outlined in (Fox *et al*., 2011; Huynh and Carruthers, 2009). For N-terminal tagging of ROCY1 in TgPRUΔKu80ΔHXGPRT a gRNA target site and PAM was identified in the region encoding the N-terminus of ROCY1 and this gRNA was cloned into pUniversal-CAS9. To generate the targeted insertion of a FLAG tag in the N-terminus, a reverse primer was designed with a homology arm to insert a single FLAG tag after the start codon in frame with residue 2 of the predicted ROCY1 protein. The forward primer amplified upstream of the HXGPRT cassette in the pGRA-HA HPT plasmid which contained the full length endogenous promoter for ROCY1, while it’s 5’ end had a homology arm that would drive homologous recombination upstream of the ROCY1 promoter in the recipient genome. This PCR product was co-transfected with the CAS9 plasmid that also encoded the N-terminus-targeting gRNA and following selection with MPA/Xanthine clones were isolated by limiting dilution. Clones were screened for FLAG-tagged ROCY1 by induced cyst formation for up to 3 days *in vitro* and staining with the M2 anti-FLAG monoclonal antibody followed by relevant secondary antibodies. Clones were also validated by western blotting using standard approaches.

### Generation of GFP luciferase parasites

TgVEG WT, TgVEGD*ROCY1,* and TgVEGD*BFD1* parasites were modified to contain GFP and click beetle luciferase using the pClickLUC-GFP plasmid (Boyle *et al*., 2007). A gRNA targeting the SNR1 gene in *T. gondii* (TgVEG_290860) was designed using the E-CRISP design tool (*Toxoplasma gondii* genome, medium setting) and incorporated into the pCRISPR_ENZ plasmid using Q5 mutagenesis and verified with Sanger sequencing (pCRISPR_SNR1). Homology arms corresponding to the 20bp sequencing flanking the Cas9 cut site were inserted to the pClickLUC-GFP plasmid (pClickLUC-GFP_SNR1-HA) using Q5 mutagenesis and verified with Sanger sequencing. Approximately 10×10^6^ TgVEG WT, TgVEGD*ROCY1,* and TgVEGD*BFD1* parasites were transfected with 25µg of the (pCRISPR_SNR1) and 25µg of the pClickLUC-GFP_SNR1-HA as described above. After ∼24 hours, the infected host cells were grown in selection media containing cDMEM and 3×10^-7^M sinefungin. After a stable population of sinefungin resistant parasites were obtain, parasites were cloned using limiting dilution and screened for expression of GFP using an Olympus IX83 inverted fluorescent microscope. To screen GFP positive parasite clones for luciferase expression, parasites were scraped, syringe lysed with a 25-and 27-gauge needle, pelleted, and resuspended in PBS at a concentration of 500,000 parasites/mL, 250,000 parasites/mL, and 125,000 parasites/mL. For each concentration of parasites, 200µL was added to each well of a black 96 well plate (100,000, 50,000, and 25,000 parasites per well) in triplicate and 50µL of d-Luciferin potassium salt was added to each well. Parasites were incubated with d-Luciferin at room temperature for 10 minutes and the luciferase signal was measure using an IVIS Lumina II *in vivo* bioluminescence imaging system. Non-luciferase expressing parasites and PBS were also used as controls for this experiment. To determine if the addition of GFP and luciferase altered tissue cyst formation, tissue cyst formation assays and immunofluorescence assays were performed as described above except parasites were only maintained in control conditions (cDMEM, pH 7.2) for 24 hours before the alkaline pH induction was performed.

### Murine TgVEG WT-GFP-LUC, TgVEGΔ*ROCY1*-GFP-LUC, and TgVEGΔ*BFD1*-GFP-LUC infections

For *in vivo* infections, 5-week-old CBA/J female mice (Jackson Laboratories) were infected with 250,000 TgVEG WT-GFP-LUC, TgVEGD*ROCY1*-GFP-LUC, and TgVEGD*BFD1*-GFP-LUC parasites (N=6) in 200 µL of PBS via intraperitoneal injection. Mice were imaged ventrally using a 4 minute exposure and large binning at 3 hours post infection and on days 1-5, 7-8, 10, and 12 post infection 10 minutes following intraperitoneal injection with 200 µL of d-Luciferin potassium salt as previously described (Saeij *et al*., 2005). After 21 days of infection, mice were sacrificed and whole brains were removed. A brain homogenate was prepared by passing whole brains through a 100 µm cell strainer using 5mLs of PBS. An additional 20 mLs of PBS was added to the homogenate and it was pelleted by spinning at 1,000 xg for 5 minutes. The supernatant was discarded, and the pellet was resuspending in 1mL of PBS, which produced a final volume of 1.2 mLs of brain homogenate.

To quantify the number of parasite genomes in the brain homogenate, genomic DNA was extracted from 100 µL of brain homogenate using the GeneJET genomic DNA isolation kit according to manufacturer’s instructions. qPCR was used to quantify the total number of genomes using primers targeting the *T. gondii* B1 gene and primers targeting mouse GAPDH as a control gene. All reactions were performed in duplicate using a QuantStudio 3 Real-Time PCR System. Samples were analyzed in 10 µL reactions consisting of 5 µL of 2X SYBR Green 2X master mix, 1 µL of 5 µM forward and 5 µM reverse primers, 2 µL of ddH2O, and 2 µL of genomic DNA. Cycling and melt curves were performed as previously (Sokol *et al*., 2018). To determine the total number of parasite genomes per brain, a standard curve of known parasite numbers was also performed using *T. gondii* B1 primers. Statistical significance was determine using DC_T_ (TgB1 C_T_ - mGAPDH C_T)_ values.

To quantify *in vivo* tissue cyst formation, 100 µL of brain homogenate was fixed with 900 µL of 4% paraformaldehyde in PBS for 20 minutes. Fixed samples were pelleted at 5,200xg for 5 minutes, washed with 1mL of PBS, resuspended in 1 mL of PBS, and stored at 4 degrees C. Fixed brain homogenate was stained with Rhodamine-labeled *Dolichos Bifluorus* Agglutinin (1:150) overnight at 4 degrees C with rotation. The following day, samples were pelleted (5,200 xg for 5 minutes) washed with 1 mL of PBS, pelleted, and resuspending in 1mL of PBS. For each sample, 50µL of stained homogenate (total of 300 µL) was added to 6 wells of a flat-bottomed 96 well plate and tissue cysts were blindly counted using an Olympus IX83 fluorescent microscope using the 10X objective. Cyst burdens were calculated by multiplying the total number of tissue cyst observed in the 300 µL of brain homogenate by the dilution factor.

### Reactivation of chronic murine infections with TgVEG WT, TgVEGΔ*ROCY1,* and TgVEGΔ*BFD1*

For *in vivo* infections, 6-week-old CBA/J female mice (Jackson Laboratories) were infected with 250,000 TgVEG WT-GFP-LUC, TgVEGΔ*ROCY1*-GFP-LUC, and TgVEGΔ*BFD1*-GFP-LUC parasites (N=12) in 200 µL of PBS via intraperitoneal injection. For each parasite, 6 mice were assigned to either the control group or the reactivation group. The reactivation group was given dexamethasone (20 mg/L) in their drinking water beginning at day 30 post infection. Mice were imaged (as described above) prior to dexamethasone treatment (0 days post dexamethasone) on days 4, 6, 7, 8, 10, 12, and 14 post-dexamethasone treatment. Peritoneal cells were collected from moribund mice and used to infect HFFs grown in T-25s. The mice belonging to the control group were sacrificed at 9 weeks post infection. Whole brains were removed, and a brain homogenate was generated as described above.

To quantify *in vivo* tissue cyst formation at 9 weeks post infection, 300 µL of brain homogenate was fixed with 1,200 µL of 4% paraformaldehyde in PBS for 20 minutes. Fixed samples were stained with Rhodamine-labeled *Dolichos bifluorus* Agglutinin as described above. For each sample, the total number of tissue cysts present in 600 µL was blindly counted by adding 50 µL of stained homogenate to 12 wells in flat-bottomed 96 well plate. Total cyst burdens were calculated by multiplying the total number of tissue cyst observed by the dilution factor.

To quantify the number of parasite genomes in the brain homogenate at 9 weeks post infection, genomic DNA was extracted from 100 µL of brain homogenate as described above. qPCR was also performed as described above.

---

**Figure S1.**
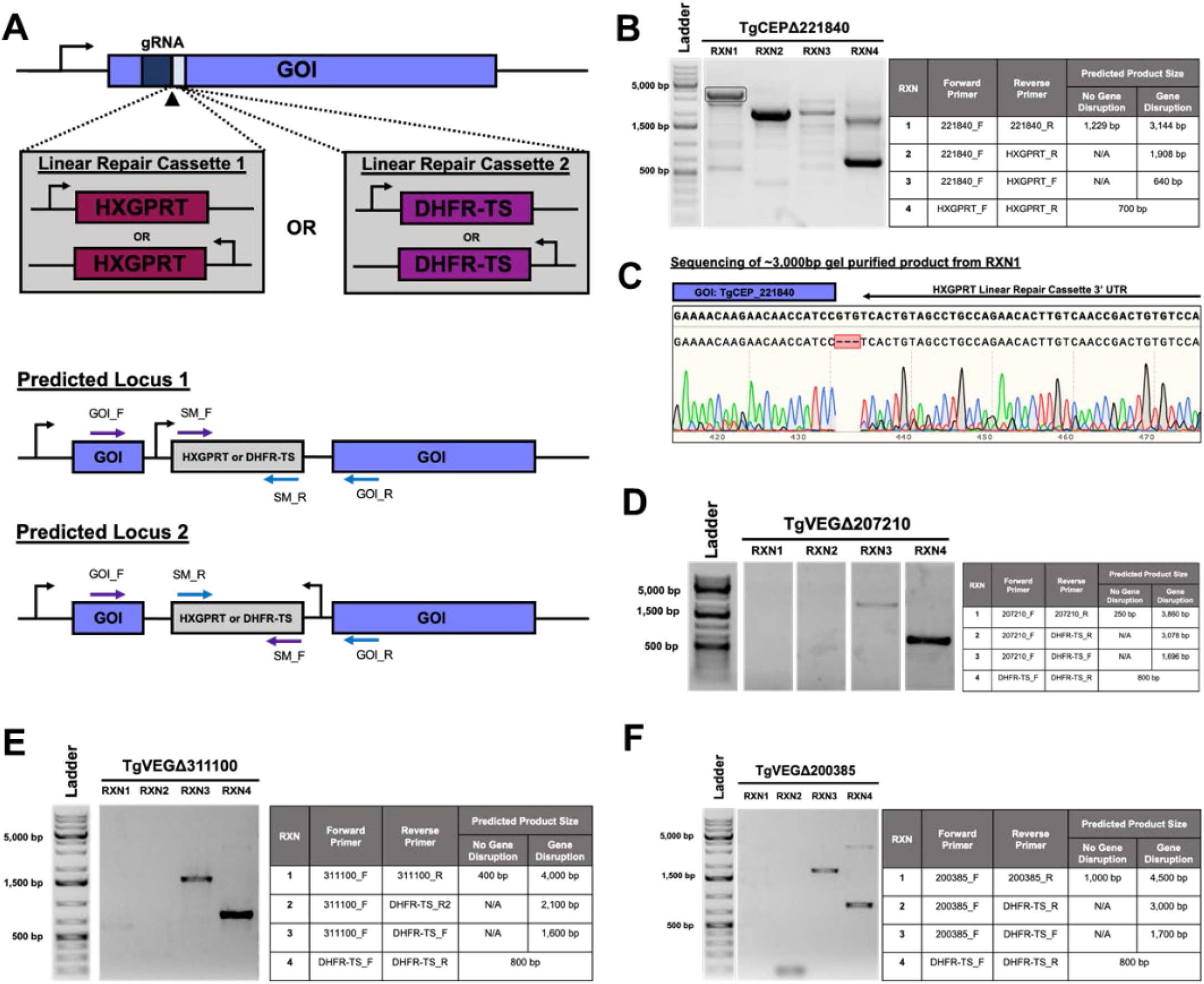
Generation of knockout of high priority candidate genes using CRISPR/Cas9. A) Schematic of CRISPR/Cas9 knockout and repair strategy and diagram of predicted loci of gene of interest (GOI) after incorporation of linear repair cassettes of selectable markers (SM) HXGPRT or DHFR-TS. Arrows indicate general location of primers. B) Gel image and PCR reactions used to identify disruption of the TgCEP_221840 locus. C) Sanger sequencing data of of gel purified ∼3,000 bp band from RXN1 sequenced with 221840_F primer. D) Gel image and PCR reactions used to identify disruption of the TgVEG_207210 locus. E) Gel image and PCR reactions used to identify disruption of the TgVEG_311100 (*ROCY1*) locus. F) Gel image and PCR reactions used to identify disruption of the TgVEG_200385 (*BFD1*) locus.

**Figure S2.**
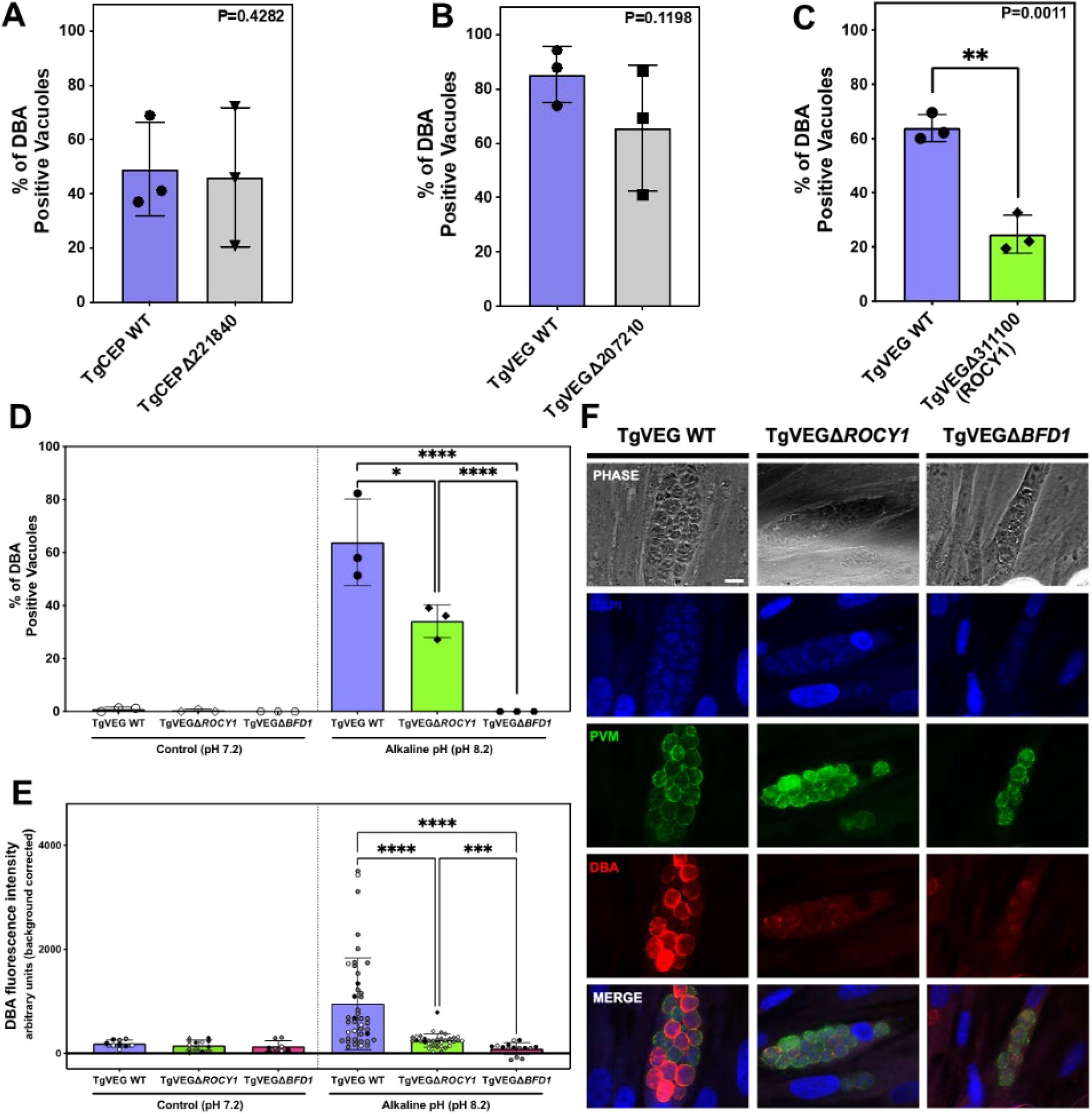
Disruption of the *ROCY1* locus impairs *in vitro* tissue cyst formation in *T. gondii* VEG parasites. A-C) Quantification of tissue cyst formation in knockout parasites A) TgCEPΔ221840 B) TgVEGΔ207210 C)TgVEGΔ311100 (*ROCY1*) and passaged matched WT strains after 48 hours of alkaline pH-induced stress. The mean and standard deviation (N=3 biological replicates) of the % DBA-positive cysts observed in 15 parasite containing fields of view is plotted. Statistical significance was determined using a Student’s one-tailed t-test on arcsine transformed percentage data. D) Quantification of tissue cyst formation in TgVEGΔ*ROCY1,* TgVEGΔ*BFD1,* and passaged matched TgVEG WT strains after growth in control conditions or after 48 hours of alkaline pH induced stress (pH 8.2). The mean and standard deviation (N=3 biological replicates) of the percentage of DBA positive vacuoles observed in 15 parasite containing fields of view is plotted. Statistical significance was determined for the alkaline pH condition using ANOVA with Tukey’s multiple comparisons tests on arcsine transformed percentage data. *p=0.025, ****p <0.0001 E) Quantification of fluorescence intensity TgVEGΔ*ROCY1* and TgVEGΔ*BFD1* from E. The mean and standard deviation of the fluorescence intensity (background corrected) is plotted. Each point represents a single vacuole. Black, gray, and white symbols indicates from which replicate (N=3) the intensity measurement was derived. Statistical significance was determined using a Brown-Forsythe and Welch ANOVA test with Dunn’s multiple comparisons.***p=0.0006 ****p <0.0001 G) Representative images of vacuoles exposed to alkaline pH stress for 48 hours. DNA was stained with DAPI (blue), the parasitophorous vacuole membrane (PVM) was stain with an anti-MAF1b mouse polyclonal antibody (green), and tissue cysts were stained with rhodamine-labeled *Dohlichos Biflorus* Agglutinin (DBA) (red). Scale bar represents 10 µm.

**Figure S3:**
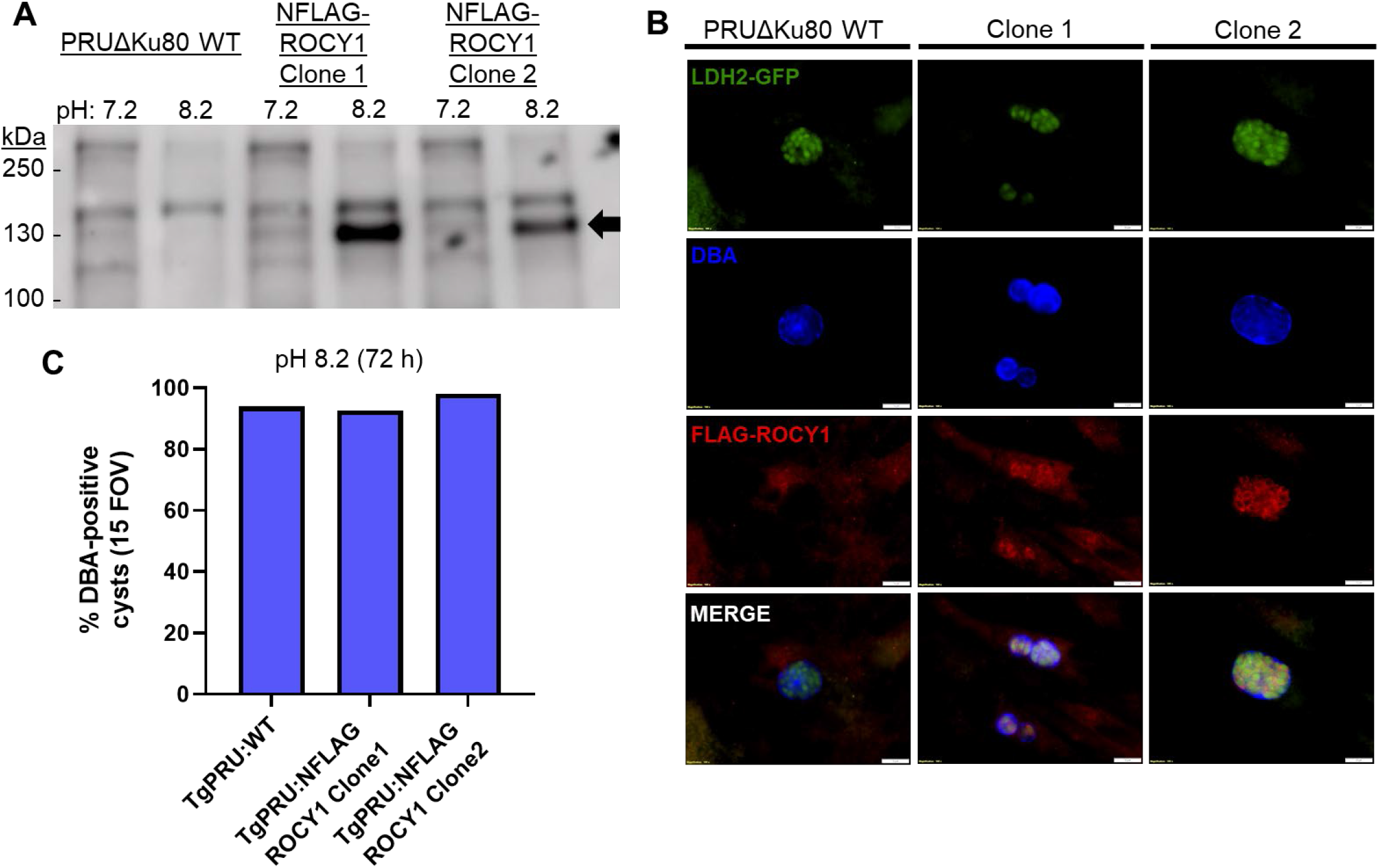
An single N-terminal FLAG tag allows for quantification and localization of the endogenous *T. gondii* ROCY1 gene product and does not impact stress-induced cyst formation. A) Indicated parasites were grown under control or high pH conditions for 3 days and then harvested for western blotting. A band with an apparent molecular weight of ∼130 KDa can be seen to increase in high pH conditions only in the FLAG-tagged clones (arrow). B) Representative images for Pru Δku80 WT, NFLAG ROCY1 Δku80 Clone 1, and NFLAG ROCY1 Δku80 Clone 2 after 3 days after switch conditions (low CO2, high pH). Parasites endogenously express GFP driven by the LDH2 promoter, cysts were stained with biotinylated DBA and ROCY1 was localized using anti-FLAG antibodies. Detectable expression of NFLAG-ROCY1 can be seen localizing to perinuclear puncta, similar to that observed for C-terminally tagged parasites (Figure 2). C) Quantification of cyst formation capacity in TgPRU:WT and the two N-terminally FLAG-tagged clones, showing that the N-terminal FLAG tag has no detectable impact on cyst formation.

**Figure S4:**
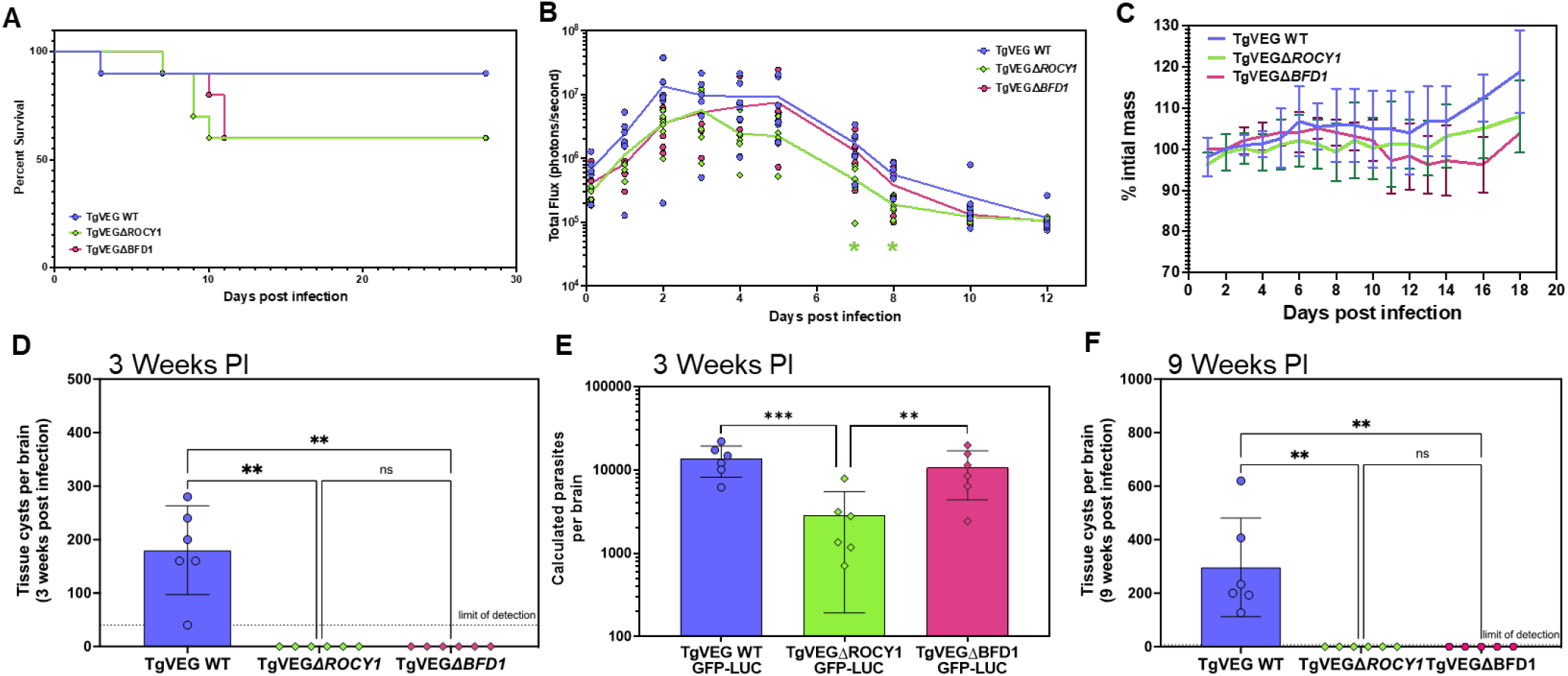
A) Mouse mortality over the course of acute infection with TgVEG wild type or TgVEGΔROCY1 or TgVEGΔBFD1 parasites. While mortality rates were higher in mice infected with the knockout parasites this difference was not statistically significant. B,C) In vivo BLI data quantifying parasite burden (B) or mouse weight loss (C) over the course of the acute phase of infection with the indicated parasite strains. This experiment is a nearly identical repeat of that outlined above in Figure 7. D,F) Quantification of cysts per brain in mice infected with the indicated strains at 3 (D) and 9 (F) weeks post-infection. E) Quantitative PCR based estimate of the number of parasites per brain at 3 weeks post-infection. While all 3 strains could be detected in the brain of the infected mice, TgVEGΔROCY1-infected mice had significantly lower numbers of parasite genome equivalents. **:P<0.01; ***: P<0.001.

**Figure S5.**
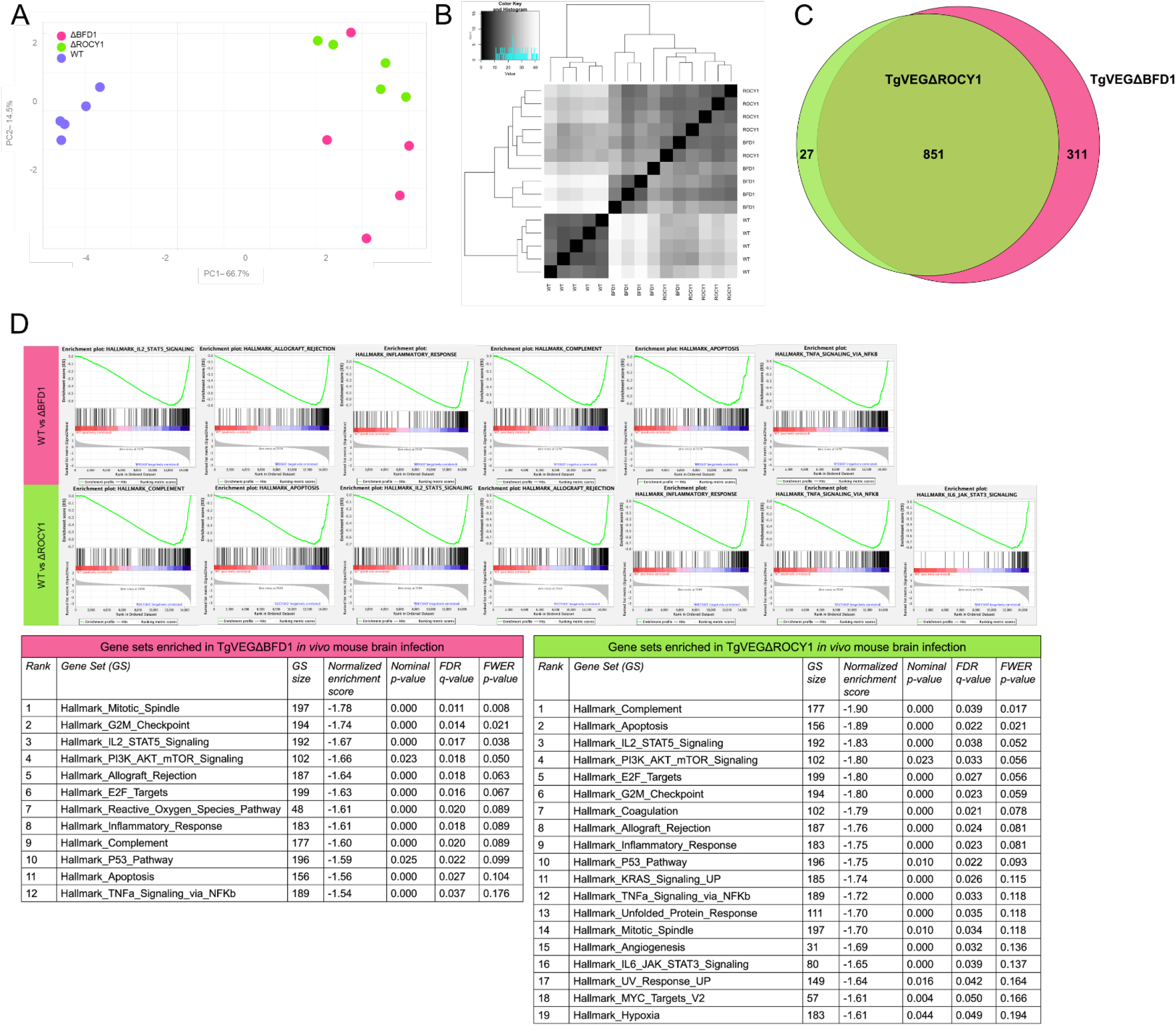
TgVEGΔBFD1 and TgVEGΔROCY1 infection initiates more inflammatory signaling in mouse brains as compared to TgVEG WT at 4 weeks post infection. A) Principal components (PC) 1 and 2 of sequenced RNA isolated 4 weeks post infection from mouse brains from *in vivo* TgVEGΔBFD1, TgVEGΔROCY1, and TgVEG WT infections. B) Unsupervised clustering of sequenced RNA isolated from mouse brains from *in vivo* TgVEGΔBFD1, TgVEGΔROCY1, and TgVEG WT infections. C) Venn diagram of genes with significant differences in transcript abundance in knockout versus WT *in vivo* infections (TgVEGΔROCY1 versus WT, and, TgVEGΔBFD1 versus WT, respectively) For both, log2 fold change < −1, adjusted p < 0.05. D) Gene set enrichment analysis indicating enriched hallmark datasets between TgVEGΔBFD1, TgVEGΔROCY1, and TgVEG WT *in vivo* mouse infections. Enrichment plots displayed indicate immunologically-relevant hits, rank-ordered by normalized enrichment score (NES), FDR q-value < 0.05. Tables show the full GSEA report of enriched hallmark pathways.

**Figure S6:**
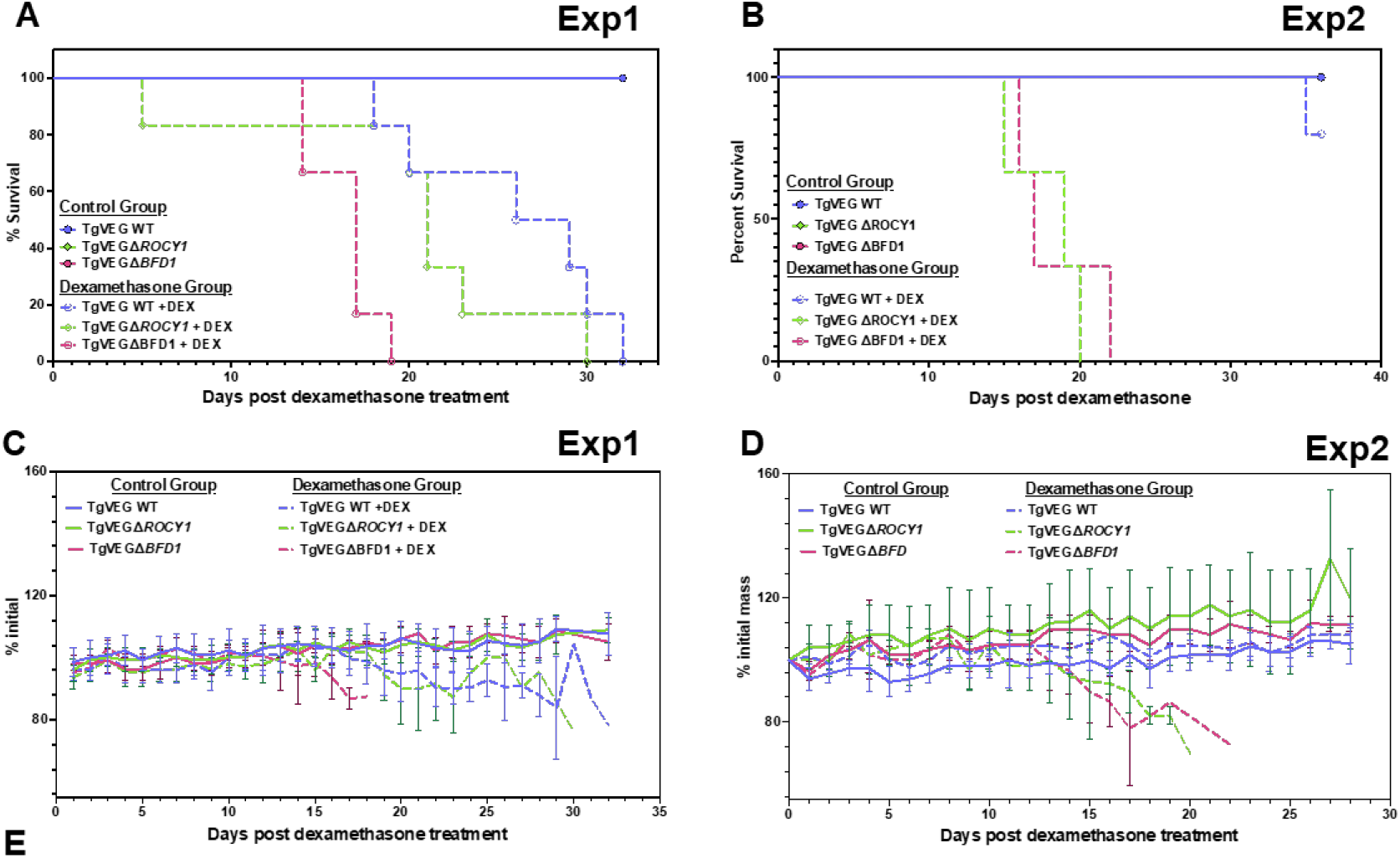
Mortality (A,B) and mouse weight loss (C,D) data for the reactivation experiments described in Figure 9B-C. Mice harboring prior infections with either ΔROCY1 or ΔBFD1 parasites succumbed to the reactivated infection significantly sooner than those infected with T. gondii VEG wild type parasites (P<0.05 for all data).

